# Individual variations in reward-seeking adaptability emerge among isogenic mice living in a micro-society and predict their response to nicotine

**DOI:** 10.1101/2023.10.28.564522

**Authors:** SL. Fayad, LM. Reynolds, N. Torquet, S. Tolu, S. Mondoloni, C. Nguyen, R. Justo, S. Didienne, N. Debray, C. Viollet, L. Raynaud, Y. Layadi, B. Hannesse, A.-M. Capaz, T. Topilko, N. Renier, A. Mourot, F. Marti, Faure Ph

## Abstract

Individual animals differ in their traits and preferences, which shape their social interactions, survival, and susceptibility to disease, including addiction. Nicotine use is highly heterogenous, and has been linked to the expression of personality traits. Although these relationships are well-documented, we have limited understanding of the neurophysiological mechanisms that give rise to distinct personalities and their connection to nicotine susceptibility. To address this question, we conducted a study using a semi-natural and social environment called “Souris-City” to observe the long-term behavior of individual mice. Souris-City provided both a communal living area and a separate test area where mice engaged in a reward-seeking task isolated from their peers. Mice developed individualistic reward-seeking strategies when choosing between water and sucrose in the test compartment, which, in turn, predicted how they adapted to the introduction of nicotine as a reinforcer. Moreover, mouse profiles in isolation also extended to correlate with their behavior within the social environment, linking decision-making strategies to the expression of behavioral traits. Neurophysiological markers of adaptability within the dopamine system were apparent upon nicotine challenge, and were associated with specific profiles. Our findings suggest that environmental adaptations influence behavioral traits and sensitivity to nicotine by acting on dopaminergic reactivity in the face of nicotine exposure, potentially contributing to addiction susceptibility. These results further emphasize the importance of understanding inter-individual variability in behavior to gain insight into the mechanisms of decision making and addiction.

## Introduction

Individuals differ not only in the way they adapt to an environment, but also in their susceptibility to disease. This is particularly evident in the context of addiction, as not all individuals will develop drug abuse despite equal exposure to a given psychoactive substance^1^. Each member within a population expresses a unique repertoire of consistent behaviors, distinct from that of its conspecifics. This inter-individual variability defines personality, a concept commonly used for humans, but which also holds true for mice^2–5^. However, in animal research, inter-individual variability in behavior has largely been considered as unwanted noise or experimental confound, and thus ignored. As a consequence, the mechanisms underlying the emergence of distinct personalities, and their role in determining vulnerability to drug use, are poorly understood.

Capturing and interpreting inter-individual variability in animal experiments can be challenging ^6,7^. Not only does individual behavior sampling require a longitudinal and complex quantification process, but the experimental design is also crucial to approach naturalistic conditions, i.e how animals living in a micro-society deal with complex and ethologically valid decision-making problems defined by the particular condition of their habitat. For instance, resource foraging is a very important aspect of animal life, and represents one of the fundamental mechanisms studied in neuroeconomics^8–10^. In most operant conditioning experiments, animals work for a food or drink ration during a time-limited daily session, before receiving complementing resources. In such an approach, individuals will typically consume according to a daily amount or body weight target, which requires pre-session restrictions as an explicit motivating operation. In addition, the timing of the meal is experimentally defined, further occulting the animal’s will to seek for resources. Instead, we studied resource foraging in a behavioral paradigm based on closed-economy^11–13^, where food and liquids are always present in a continuous experiment, 24 hours a day, and animals live in enriched groups (∼10 mice) representing a ‘micro-society’. Rodents are indeed social animals; this is aptly evidenced by their repertoire of various interactive behaviors - such as physical contact, vocal communication, aggression, social recognition - that can be considered as hallmarks of sociability, an important personality trait ^14^. Importantly, when mice live in micro-societies within a closed and enriched naturalistic environment, strong and stable inter-individual variability in behavior emerges, even among isogenic animals ^6,7^.

Decision-making behavior and strategy are known to be markedly individualistic. Behavioral trait components of decision making, such as impulsivity, exploration, or novelty seeking, are thought to predict vulnerability to drugs of abuse ^15,16^. While these personality traits have indeed been linked with smoking and addiction to nicotine in humans and animal models, certain traits, like impulsivity or sensation-seeking, have been more strongly associated with initial nicotine sensitivity^17–19^, suggesting that they are a measure of vulnerability to nicotine. However, whether the processes leading to nicotine addiction and the mechanisms of decision-making share mechanistic underpinnings remains elusive. Altered dopamine circuit function is a promising mechanistic candidate ^20,21^, as dopaminergic signaling is implicated in decision-making, in social behaviors, and in nicotine addiction, where the initial stage critically involves activation of mesolimbic dopamine neurons^22–24^. As such, investigating variations in nicotinic control over the DA system represents a particularly promising avenue for linking personality, decision-making, and vulnerability to nicotine. Nicotine initiates reinforcement by increasing the firing rate and bursting activity of DA neurons through direct actions on nicotinic acetylcholine receptors (nAChRs), a family of pentameric ligand-gated ion channels with twelve different types of subunits (α and β) expressed in the mammalian brain. It has been shown that the transition between tonic and phasic activity of DA neurons induced by nicotine is essential for the reinforcement^25,26^, and that the expression of nicotine-sensitive nAChR subtypes in the VTA is necessary for both the cellular and behavioral effects of nicotine^25,27–29^. Under nicotine-free conditions, nAChRs in the VTA are also key modulators of DA activity through basal cholinergic signaling, and they regulate specific aspects of reward-seeking behaviors, in particular exploration and reaction to uncertainty^30,31^. Environmental manipulations that alter nAChR-mediated control of DA neurons may therefore lead to changes in downstream behaviors. For example, nicotine exposure modifies exploratory behavior^32,33^, by increasing, in mice, DA neuron activity and biasing individual strategies toward reduced exploration ^34^. In addition, specific social contexts (i.e., repeated aggression) have been shown to induce a marked remodeling of the dopaminergic and nicotinic system, leading to increased VTA DA neuron activity ^35,36^, social aversion, and modified nicotine response^37^. This crosstalk at the level of the DA system between responses to drugs and modifications of decision-making, could explain the observed correlation between novelty seeking and susceptibility to nicotine.

Here, we aim to demonstrate whether the way individuals adapt to their environment can differentially modify nicotinic modulation of DA networks among individuals, and, consequently, the sensitivity of mice to nicotine, a critical element that may define susceptibility^38,39^. For that purpose, we used a habitat called “Souris-City” that combines a large social environment where mice live together with a modular testing platform where animals individually perform cognitive tests. In this environment, mice have individual access to water by performing a specific task in a T-maze, while social, circadian, and cognitive behaviors are continuously monitored over time using multiple sensors^7^.

## Results

### Souris-City: continuous tracking of individual mice living within a micro-society

Souris-City is a semi-naturalistic environment composed of a large and complex housing space in which groups of mice (N=32 groups) live together (5 to 10 male mice per group, mean = 8.8) for extended periods of time (1–3 months) and are able to express sophisticated social and non-social behaviors (Figure 1A). The environment includes a test-area (individual zone), separated from the main environment (social zone) by a gate which selectively controls the passage of mice, one at a time, based on a Radio Frequency Identification (RFID). Mice can thus perform a self-initiated cognitive task individually, spontaneously, and isolated from their cage mates. The test area consists of a T-maze leading to two drinking areas at the end of the left and right arms. Thus Souris-City associates a zone for individual liquid consumption and a social zone (the main cage) where food is always available. The experimental paradigm involves several consecutive periods with modified rules regarding access to and the nature of the liquids (Figure 1B). During a one-week habituation period, mice explore Souris-City with free access to the T-maze. The gate is always open, so several mice can access the T-maze simultaneously, and water is delivered from both sides. In the second period (WW, mean duration = 9.5 days, n=281 mice), mice continue to have water on both sides of the T-maze, but its access is now restricted by the gate so that mice can only enter the T-maze one at a time. Choice is restricted so that if the animal chooses one side, access to the opposite arm closes. During the subsequent weeks (WS, mean duration = 25.2 days, n=281 mice), water and a 5% sucrose solution are respectively delivered on each side of the gate-restricted T-maze, thus introducing a choice (choosing left or right) which modifies the reward associated with liquid consumption. The position of water and sucrose bottles are swapped twice a week, which allows for the mice to stabilize their choice. Overall, in 32 experiments (or groups of mice) in the water-sucrose (WS) test period, the behavior of 281 mice and more than 100 000 choices were analyzed.

**Figure 1:**
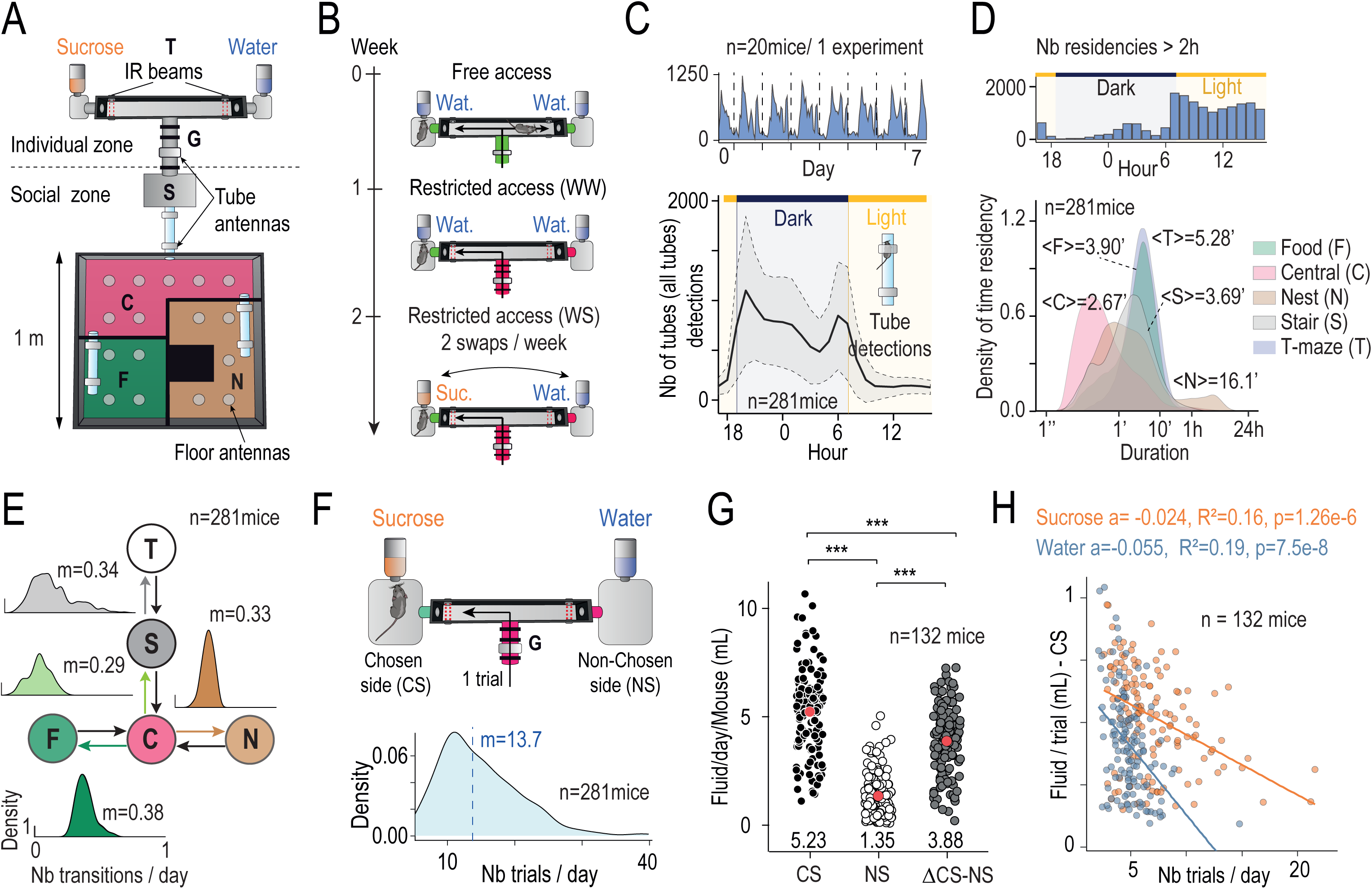
Longitudinal profiling of individual and group behavior among mouse micro-societies within Souris-City. (**A**) Souris-City is divided into two main parts: a social zone and a test zone. The social zone includes a square cage measuring 1 m × 1 m, which is further divided into four compartments: the nest (N), the food (F) area, where mice have unrestricted access to food, and the central (C) zone that serves as a hub, connecting the social compartments with a stair (S) leading to the test zone. The test zone is a T-maze, which is separated from the stair by a controlled access gate (G). Mice are tagged with RFID chips and detected using floor-mounted circular or tube-shaped RFID antennae, which connect compartments of SC to capture transitions between zones. Two infrared beams (red dashed lines) are used to detect which arm mice choose in the T-maze. **(B)** The experimental paradigm involves several consecutive sessions with modified rules regarding access to the maze and the nature of the liquids available at each arm. During the free access period (*Top*), mice are allowed unrestricted access to the T-maze for one week. The gate remains open, allowing multiple mice to enter the T-maze simultaneously, and water is delivered from both sides. In a second step (*Middle*) mice choose between water on both sides of the T-maze (WW, mean duration = 9.5 days), however access to the T-maze is restricted by the gate, and mice may only enter the T-maze one at a time. Choice is restricted so that if the animal chooses one side, access to the opposite arm is closed. Finally, water and 5% sucrose solution (*Bottom*, WS, mean duration = 25.2 days) are respectively delivered at each side of the gate-restricted T-maze, introducing a choice (choosing left or right, choosing water or sucrose). The position of the water and sucrose bottles are then swapped twice a week. (**C**) Overall activity of mice captured from their movement in Souris-City reflects their circadian rhythm. (*Top*) Tube detection events for eight consecutive days (n=20 mice, 2 group of 10 in parallel). (*Bottom*) Daily tube detection events per hours averaged for all mice (mean ± SEM, n=281) (**D**) Residency time in each sub-compartment can be captured by floor antennae. (*Top*) Histogram (bin per hour) of the number of residencies in the nest zone longer than two hours. (*Bottom*) Density of residency time in each sub-compartment (log-scale, bandwidth = 0.1), with indicated mean value. (**E**) Tube antennae provide information about the movement of mice between sub-compartments. Flow diagram of all possible transitions between sub-compartment, density graph above each transition indicates the distribution of conditional transition probability among the n=281 mice, with indicated median value. (**F**) Distribution of mean number of T-maze entries per day for n=281 mice in the SW session. Vertical dashed line indicates mean value. (**G**) Estimation of daily consumption on a subpart of the experiment (n=132 mice, see text) during SW session: Mean daily fluid change per animal distinguishing the chosen side (CS) from the non-selected side (NS) and the difference between the two (Δ) (Pairwise comparisons using Wilcoxon rank sum test with continuity correction and holm p-value adjustment correction, n= 132 mice) (**H**) Mean fluid change (chosen side) per trial per animal for trial where water (blue) or sucrose(orange) is chosen, depending on the number of trials per days (linear regression, a indicated the slope estimate, R^2^ the Adjusted R-squared and p the p-value, n= 132 mice). Data are represented as mean ± sem. ns p>0.05, ** p<0.01, *** p<0.001.

The data obtained in Souris-City are based on the tracking of animals implanted subcutaneously with RFID chips and detected by antennae placed throughout the floor and the tubes connecting the different sub-compartments of the environment (see Methods). Mice have free access to a nest sub-compartment (N), a food sub-compartment (F), a central compartment that provides access to all other compartments (C), the stair (S), and finally the T-maze (T). These subdivisions allow animal trajectories in Souris-City to be represented as a sequence of residency times within a sub-compartment and transitions between sub-compartments^31,40^. The circadian rhythm of the group emerged from the measurement of pooled activity estimated from RFID detection at the level of the transition tubes during the WS period (Figure 1C). As expected, the mice are more active (and therefore more frequently detected moving between sub-compartments) during the dark phase (7p.m. to 7a.m) than during the light phase (7a.m. to 7p.m). The time spent by mice in a given sub-compartment during the WS period varied between tens of seconds to hours, with the shortest visits, mainly found in C, corresponding to transition episodes (Figure1D *Bottom*). Time residency in the nest sub-compartment shows a bimodal distribution, with the longest occupancies observed in the environment lasting more than 2h. The distribution of long occupancy episodes, which took place mainly in the nest compartment, shows that they occurred mostly during the light period (Figure 1D, *Top*), thus interpreted as sleeping episodes. The distribution of transition probabilities from one compartment to another (Figure 1E), starting from the central position of C, shows a preponderance of transitions from C to F (median=38%) rather than towards the N (33%, Wilcoxon signed rank test p<2.2e-16) or the S (29%, Wilcoxon signed rank test p<2.2e-16). Furthermore, when animals are in S, their probability of entering the T is only 34%. This relatively low rate reflects the fact that animals enter the stair without necessarily succeeding to enter the T-maze, and then return to the main environment. Mice enter the T an average of 14.7 times per day during the WS period (Figure 1F), however the distribution is skewed with a median at 13.7 and a peak at 11 times per day and a long tail indicating that some mice can enter more than 30 times per day. In restricted access sessions, when a mouse chooses one side the access to the bottle on the other, non-chosen, side is closed, so that the mouse has access to only the bottle on the chosen side. The mouse will only be able to access the bottle on the non-chosen side if it leaves the T-maze and returns for another trial, which will reopen access to both bottles, as well as reopening access for other mice to enter. In half of the experiments (16/32 corresponding to n = 132 animals), fluid consumption was estimated for each trial (i.e., each passage of a mouse in the T-maze). By comparing the average difference in liquid change between the chosen side and the non-chosen side per day and per animal (Figure 1G, n = 132), we find a significant difference in the amount of liquid dispensed depending on the arm chosen. We thus estimate the consumption of the animals by subtracting the loss of liquid measured by the system in the non-chosen side (resulting from evaporation, noise, etc.) from the change in fluid volume measured from the bottle in the chosen side, which results in an average consumption of ∼3.9 ml per day per mouse (Figure 1G).

Consumption volume per trial depended on both the bottle content (water or sucrose) and the number of trials per day for each mouse (Figure 1H). These first analyses describe a set of average behaviors, accessible from the analysis of events captured by RFID antennae or consumption sensors. They begin to reveal an organization of behaviors with important variations in their expression depending on the individual mouse.

### Multidimensional analysis of idiosyncratic reward-seeking behavior reveals that mice adopt distinct strategies in the T-maze

In the T-maze, mice (n=281) isolated from their peers, voluntarily performed a relatively simple decision-making task: whether to make a left or right turn to access a liquid reward. In the WS test sessions, one drinking area at the end of one of the T-maze arms contains water, and the other contains a sucrose solution. Each entrance into the T-maze, and the subsequent choice of which side to access, is considered a trial (Figure 2A, *Top*). The sides of the sucrose and water bottles are swapped every 3-4 days, with each swap defining the beginning of a new session. The behavior of the mice in the T-maze was assessed by five variables that quantify the animals’ choice across different time scales throughout the entire WS experimental period. The level of global switching is estimated by the variable *Switch*, which takes all WS sessions into account and gives an overview of the probability of choosing of one side compared to the other. This probability is renormalized so that 100% corresponds to an equivalent number of visits to both sides, while 0 corresponds to an individual who visits only one side. The variables *SwWat* and *SwSuc* evaluate the choices of the animals at the trial level (i.e., going left or right). They represent the probability of switching sides if the previous choice was water or sucrose, respectively. Finally, the *Pref* and *SideBias* variables assess sucrose preference (probability of sucrose choice) and side bias (probability of choices on one side) by comparing the choices between each session (i.e., whether the sucrose is on the left or right). Despite some recurring patterns, there are considerable variations in these parameters between mice (Figure 2A, *Bottom*).

**Figure 2:**
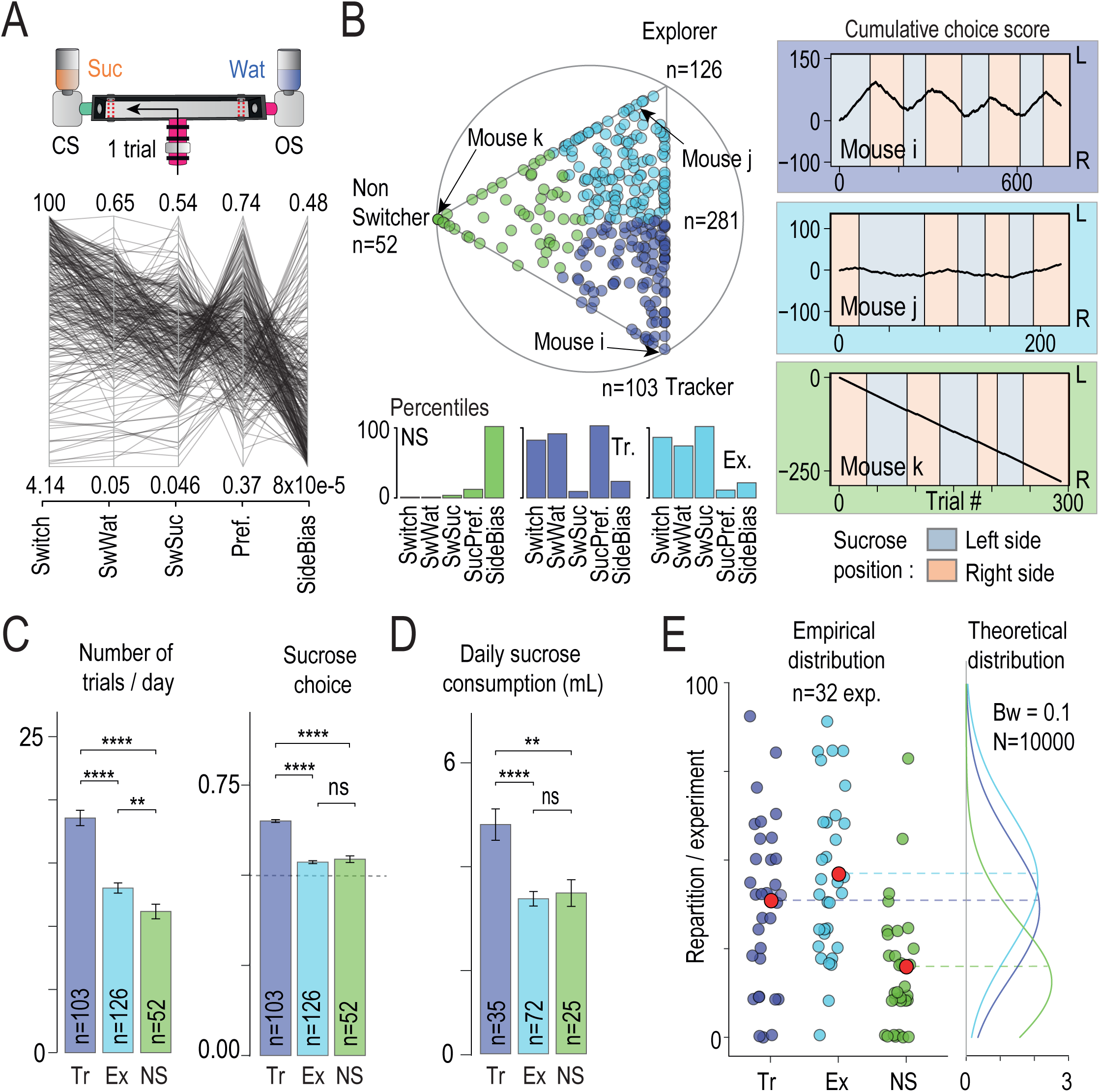
Mice exhibit inter-individual differences in choice strategies in the T-maze. **(A)** *Top*: one trial is considered to be one choice between left or right side in the T-maze. *Bottom*: Value of the five parameters that describe mice sequence of choice in the T-maze during SW sessions (n=281 mice): the level of global switching (*Switch*), the probability of switching sides if the previous choice was water (*SwWat*) or sucrose (*SwSuc),* the preference (*Pref*) and side bias (*SideBias*) on each session. **(B)** Archetypal analysis of the choice strategies based on the 5-dimensional data space. *Top*: Visualization of the α coefficients using a ternary plot. Each point represents the projection of an individual (n=281 mice) onto the plane defined by a triangle where the three apices represent the three archetypes: Tracker (Tr, purple), Explorer (Ex, blue) and non-Switcher (NS, green). Points are color-coded according to their proximity to the archetypes. *Bottom*: Histograms showing the three archetypes’ percentiles for each choice parameter. *Right*: Examples of three sequences of choice made by 3 mice close to the archetype. Sucrose position alternates across sessions between the left (light purple) and the right (light orange) side. Cumulated choices across trials are calculated with a positive (+1) or negative (-1) increment when the left or right side is chosen, respectively. The mouse i, j, k (from top to bottom corresponds respectively to a Tr., Exp. and NS profile (see their projection in the ternary plot). **(C)** Number of trials per days (*Left*) and percentage of sucrose side choice (*Right*) for the three archetypes (pairwise Wilcoxon tests with Holm correction). **(D)** Daily sucrose consumption for the three archetypes (pairwise Wilcoxon tests with Holm correction). **(E)** Repartition of archetypes per experiment showed that they are not evenly represented in each group (N=32, red dot indicated mean values, *Left*) and built theoretical densities expected for each archetype based on a random draw from mean groups sizes (Bandwidth = 0.1, n=10000, *Right*).

To better describe this variability, we used archetypal analysis, an unsupervised approach for identifying behavioral clusters. It depicts individual behavior as a continuum within an archetypal landscape defined by extreme strategies: the archetypes. The five-dimensional data set characterizing individual responses was used to identify three archetypal phenotypes. Individual data points are thus represented as linear combinations of extrema (vertex corresponding to archetypal strategies) of the data set, i.e., each mouse is represented by a triplet of α coefficients describing the archetypal composition and can be visualized with a ternary plot (Figure 2B, *Top*). The three archetypes distinguish Trackers (Tr) who track the sucrose position (Figure 2B, *Top right*, see mouse i as an example), from Explorers (Ex) who choose almost randomly between the left and right side on each trial (Figure 2B, *Middle right*, see mouse j) and Non-Switchers (NS) who choose the same side throughout the majority of the sessions (Figure 2B, *Bottom right*, see mouse k). Subsequent analysis highlights that these three profiles are distinguished not only by the choice parameters in the T-maze (used in their construction), but also by sucrose consumption and number of entries in the T-maze, reinforcing the definition of the profiles as personality-like categories. Trackers enter the T-maze frequently (i.e., high number of trials per day), whereas Non-Switchers rarely enter it (Figure 2C, *Left*). In terms of choice, Trackers go most often to the sucrose side (Figure 2C *Right*) and consume more sucrose than the others (Figure 2D). Finally, the three profiles are distributed across the different experimental groups (N = 32) tested (mean=8.8 mice per group, min = 5, max = 10) with an average proportion in a group of 36.6% for Trackers, 44.4% for Explorers and 18.5%, for Non-Switchers. The observed distribution of proportions within a group (Figure 2E, *Left*) is consistent with a random sampling (Figure 2E, *Right*) of a profile for each animal (with the corresponding probability) within group sizes similar to those obtained experimentally.

### Reward-seeking strategy in isolation correlates with behavioral trait variation in the social compartment: a proxy for mouse personality

Behavior in the T-maze is fundamentally different from behavior in the main environment. In the T-maze, animals are isolated from any direct influence from other animals and are left to make their own decisions. This is not the case for behavior in the main environment, where all behaviors are potentially subject to the consequences of social interactions. Because of these strong differences in context, we wondered whether the differences in strategy observed in the T-maze would also correspond to behavioral differences in the main environment, suggesting that mice strategies can serve as a marker of their personality across multiple levels of analysis.

The analysis of the behavior of the mice in the main environment is based on their detection by antennae located on the three transition tubes between the sub-compartments. Compared to mice with other archetypally-defined profiles, Trackers showed an increase in the average number of tube antenna detections per day (*NbD,* Figure 3A). They also have a reduced probability of transitioning from Nest to Food compartments (%*NtoF*, Figure 3B). A strong inverse correlation was observed between NbD and the %NtoF across all mice, regardless of their archetypal profile (Figure 3C). This suggests a more nuanced relationship between these variables than what can be captured by simple group statistics, and reinforces the idea that mice can be individually defined by their behavioral repertoires, indicative of a type of personality for each mouse. Because the archetypal framework defines each individual as a linear combination of profiles, it is possible to further refine this analysis by introducing the notion of distance from the archetype. The archetypal composition (i.e., given by α_k_ with k the archetype, Figure 3D) reflects this distance: its value is between 0 and 1, with 1 being if the mouse is exactly on the archetype, and 0 if it is on the opposite side of the archetypal space. We found that the %*NtoF* decreases across mice as their composition approaches the pure Tracker archetype, while their *NbD* increases (Figure 3E, *Top*). However, these two relationships are reversed for the Explorer archetype composition, such that %*NtoF* increases and *NbD* decreases as the composition of the mice approaches the pure Explorer archetype (Figure 3E, *Bottom*). These correlations reflect both profile differences and environmental constraints (i.e., the structure of the settings) on behavioral expression.

**Figure 3:**
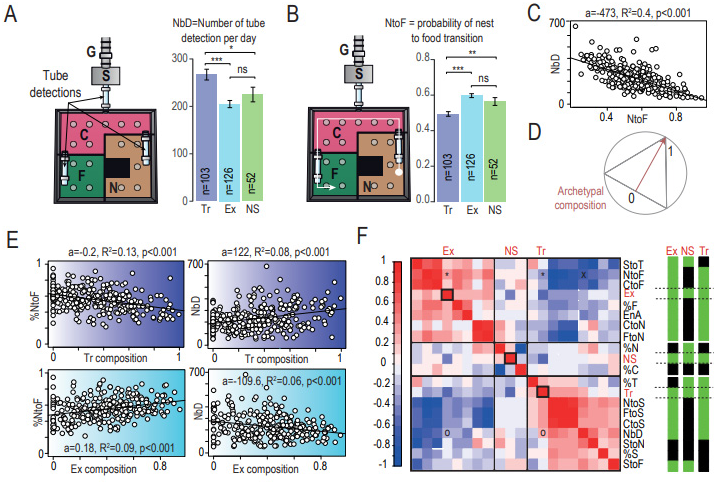
Archetypes defined by individual choices capture variation in the social cage behavior. **(A)** Activity in the main environment, estimated by the number of transitions between compartment (NbD), for the three archetypes (pairwise Wilcoxon tests with Holm correction). **(B)** Probability of nest to food transition (NtoF) for the three archetypes (pairwise Wilcoxon tests with Holm correction). **(C)** Correlation between NtoF and NbD. **(D)** Principle of archetypal composition measurement: the archetypal composition (i.e., given by α_k_ with k the archetype) would be equal to 1 if the mouse is exactly at the point of the archetype, and 0 if it is on the opposite side. **(E)** Correlation (linear regression, a indicating the slope estimate, R^2^ the Adjusted R-squared and p the p-value) between Tracker (Tr) composition and pNtoF (*Left*) and NbD (*Right*) respectively (*Top*), and between Explorer (Ex) composition and pNtoF (*Left*) and NbD (*Right*) respectively (*Bottom*). **(F)** *Left*: Correlation matrix of main environment variables and archetypal profile. *Right*: p-value for correlations. Green: p<0.05 Black: p>0.05. Variables: Activity Levels: Number of Detections (NbD), Entropy (EnA) ; Probability of Transitions: Stair to T-maze (StoT), Nest to Food (NtoF), Center to Food (CtoF), Center to Nest (CtoN), Food to Nest (FtoN), Nest to Stair (NtoS), Food to Stair (FtoS), Center to Stair (CtoS), Stair to Nest (StoN), Stair to Food (StoF) ; Occupancy : percent time in Food compartment (%F), percent time in Nest compartment (%N), percent time in Center compartment (%C), percent time in T-Maze compartment (%T), percent time in Stair compartment (%S); Archetypes : Explorer (Ex), Non-Switcher (NS), Tracker (Tr). * and ° indicates correlation between Tr and Ex composition with pNtoF and NbD shown in (E). X indicates correlation shown in C.

Therefore, we systematically analyzed the linear correlation between an individual’s archetypal composition and specific behaviors in the main environment (Figure 3F, *Right*). Three categories of variables were used to describe activity levels (Figure 1C), compartment occupancy (Figure 1D), and transitions, respectively (Figure 1E). Robust correlations were found between the variables describing these categories and the archetypal compositions, with the pattern of these correlations also discriminating between archetypal profiles (Figure 3F, *Left*). Interestingly, Explorer and Tracker archetypes often exhibit correlations with the same behaviors; however, these correlations are inversely related. This suggests that Explorer and Tracker could be construed as contrasting profiles within the primary environment. Tracker composition was positively correlated with parameters indicating a preference for the food or the nest compartment over the stair compartment (Supp Figure 3). For Explorer, the opposite is observed: these animals go more to the T and are overall more active (i.e more detected at the level of tube transition antennae, Supp Figure 3). Non-Switchers display a profile that is markedly distinct from the other two profiles (Figure 3F, *Right*). They are characterized by a preference for the nest compartment (Supp Figure 3).

Overall, these analyses suggest that individual mouse profiles extend beyond variations solely within reward seeking strategies in the T-maze to also encompass differences in activity within the main compartment. These relationships between individualistic strategy development and trait expression can be considered as a foundation of mice “personalities”, which emerge as adaptive responses to their complex environment.

### Distinct reward-seeking profiles are defined by individual differences in learning rate and sensitivity to value

Because decision making strategy is a good marker for personality profiles of mice in Souris-City, we next aimed to decompose the latent variables that individual mice use to define their strategy and relate these parameters to drug-taking vulnerability. The decision-making process of an individual mouse in the T-maze can be seen as a series of binary choices between going left or right in an unpredictable environment. It is assumed that the animal learns the value assigned to each option (left or right) and that it adapts to the change (every 3-4 days) in the position of the rewards (sucrose or water). In this context, we fit each individual’s choice data with a standard reinforcement learning model ^41^, which uses the sequence of choices to estimate the expected value of each option for each trial. The value of the chosen option (V_L_ or V_R_ for left and right side respectively) was updated after each trial using a reward prediction error rule (see methods) and a learning rate α that sets how rapidly the estimate of expected value is updated on each trial. Given expected values for both options, the probability of choosing the right option P_R_(t) is computed using a SoftMax rule with two parameters: the inverse temperature parameter β which represents the sensitivity to the difference of values and a choice perseveration parameter χ that captures short-term tendencies (previous choice) to perseverate or alternate (when positive or negative, respectively, Figure 4A *Left*). This propensity to alternate is independent of the reward history ^42^, and thus does not depend on ΔV. We fitted the choice data of each mouse with this model and obtained triplet of latent variable values (α, β and χ, Figure 4A *Right*) for each individual (see methods). The three archetypes extracted from the sequence of choice corresponded to different combinations of α, β and χ (Figure 4B). These parameters also correlate with the number of trials in the T-maze (Supp. Figure 4A, α : p = 0.002; R^2^ = 0.03 ; β : p = 7e^-8^, R^2^ = 0.1; χ : p = 2e^-5^, R^2^ = 0.06), indicating that these latent variables also capture information that are not directly linked to the decision process. Finally, when we use this model to simulate data under the constraints of experimental trial sequences and rewards (water/sucrose, right or left), we can differentiate the same three types of profiles that we find in the experimental data (Figure 4C). One question, however, is whether the estimated latent variables (Figure 4A) or the dynamics of the choices (Figure 4C) simply reflect a difference in the number of trials, as individuals who entered the T-Maze the least were indeed less likely to find the sucrose. To test this hypothesis and to decorrelate our results from a possible difference due to a variation in the number of trials, we modeled the behavioral profile from the latent variables with 6 alternating sessions (sucrose/water left or right) of 50 trials each. Our model with three latent variables (α, β, and χ) explained the phenotypic variables *(Switch, SwWat, SwSuc, Pref, and Side Bias*) very well, regardless of the number of trials (Supp. Figure 4B).

**Figure 4:**
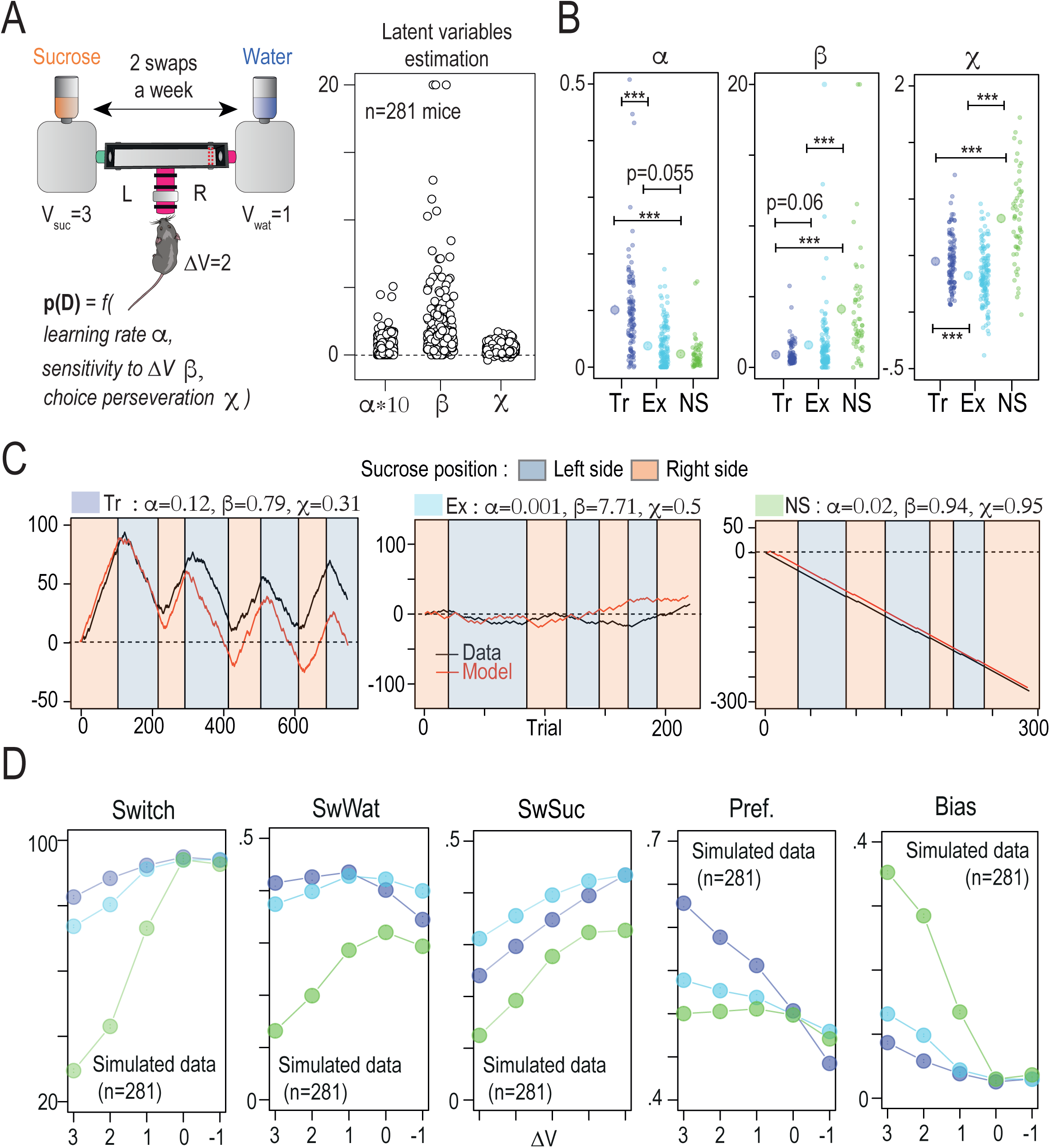
Computational modeling suggests that decision and learning parameters differ between the three archetypes. **(A)** *Right*: Principle of the reinforcement learning and SoftMax model, with three Latent variables α (the learning rate), β (inverse temperature or sensitivity to the difference of values ΔV) and χ (the choice perseveration). The theoretical value of water and sucrose are set to 3 and 1 respectively. *Left*: Estimated values of β, χ and α for n=281 mice. **(B)** Latent variables according to the Tr, Ex and NS archetype (right mean ± sem, Wilcoxon tests with Holm correction, left: individual value per mice). **(C)** The model recapitulates the profiles drawn from experimental data (same exemple as in figure 2B) when fitted with individual triplets values for the latent variables of each individual of a specific archetype. **(D)** Comparison of the mean of five variables (*Switch*, *SwWat*, *SwSuc, Pref*, *SideBias*) for Tr., Ex. and NS archetype obtained for five differences in the value (ΔV) associated with the choice (6 sessions of 50 choices, simulated with fitted values of β, χ and α).

Having demonstrated the validity of our model we next questioned the relative importance of each latent variable in explaining the observed results by designing an attribution study in which each of the latent variables (α, β or χ) is manipulated independently from the other. We then simulated the data and compared the results obtained using i) the latent variable values estimated for each mouse associated with each profile, ii) by randomly picking latent variable triplets from the set of estimated values, and iii) by using two estimated values for each individual and randomly picking the third from the set of possible values obtained (Supp. Figure 4C). This analysis reveals that the χ parameter plays a minimal role in the differentiation of the profiles. In other words, this particular parameter does not contribute significantly to the observed variations within the profiles. Overall, the Tracker archetype is associated with a high α value and low β, consistent with individuals who are able to quickly update their value representation and thus favor sucrose tracking. From one session to the next, they follow the sucrose and switch to the other side. The Explorer archetype was characterized by an intermediate α and β, and thus an important level of switching from one trial to the other. In contrast, the Non-Switchers are associated with low α and high β, a combination that favors profiles in which the animals remain mainly on one side, in particular the side with the highest initial value representation (this representation is updated slowly due to a low α). Indeed, the majority of Non-Switchers showed a bias for the side on which they first encountered the sucrose (34/52) or the side where they found sucrose for the majority of their first choices and (2/52) (Supp. Figure 4D). The other Non-Switcher mice (16/52) choose the side corresponding to their preference during the WW session.

### Reward-seeking strategy defines adaptability to changes in reward value and predicts nicotine choice

We next asked if an individual’s profile, defined as a combination of their latent variables, could predict how they would adapt to other choice situations. In a binary choice, such as in the T-maze, a major external variable is how the value between the two options differs. We thus simulated the theoretical response of the different profiles, namely Tracker, Explorer, and Non-Switcher, to changes in reward values difference (ΔV) in the T-maze. Variation in the five choice variables revealed different adaptations to ΔV depending on the profile (Figure 4D). The Ex-archetype is characterized by little adaptation of their choice with ΔV, whereas Tr and NS-archetypes show strong adaptation. Trackers show a clear side preference for the higher value, and this preference increases with the difference in reward value. Finally, the most surprising result comes from the Non-Switcher profile. Their side bias is clearly not independent of ΔV, but becomes more important as the value difference increases: the higher the ΔV, the more the mice stick to one side of the T-maze. This simulation thus suggests that our three profiles adapt completely differently to changes in reward in the T-maze.

We tested this hypothesis by conducting an experiment where we modulate ΔV between two consecutive series of sessions in mice. The first round featured a significant difference in value between the two choices (sucrose sessions), whereas the second round had a purportedly smaller contrast (nicotine session). Hence, a subgroup (n = 74) of the 281 mice was subjected to a period (SaN) of choice between nicotine (100 μg/mL) plus 2% saccharin (Nic) versus 2% saccharin alone (Sac) in the T-maze, following the prior water versus sucrose period (WS) (Figure 5A). The mouse behavior during the WS session showed a typical distribution of mice among the three profiles in the archetypal space (Figure 5B). During the SaN period, there was a global decrease in the number of successful choices (with nicotine considered the successful choice), but no significant change in the number of trials per day (Figure 5C). Additionally, the consumption pattern changed, as mice consumed more sucrose than water in the WS period, whereas they consumed as much saccharin as nicotine in the SaN period (Figure 5D), suggesting that the ΔV between nicotine+saccharin and saccharin alone is smaller than the ΔV between water and sucrose. Next, we examined changes in the five variables (Switch, SwWat, SwSuc, Pref, and Side Bias) that describe the choice behavior in the T-maze across the two periods (Figure 5E). Consistent with the simulation predictions, there were no modifications in these five variables for the Explorer between WS and SaN periods, suggesting a minimal adaptation to the change in ΔV. In contrast, significant adaptations were observed in the Trackers and Non-Switchers profiles. The Trackers displayed a decrease in preference (Suc/Nic Pref) and switch behavior after choosing the saccharin side (swWat/Sac), but an increase in switch behavior after choosing the nicotine side (swSuc/Nic). On the other hand, the Non-Switchers primarily showed an increase in switch behavior after receiving the reward (swSuc/Nic). In the SaN period, Trackers and Explorers were similar for all variables, and different from the Non-Switchers. The results demonstrate that the model and the latent variables effectively capture the information needed to model the behavior of mice under different reward values, depending on their profile. In addition, the model accurately predicts the behavioral response to the choice between nicotine+saccharin and saccharin alone, a choice with a smaller ΔV in comparison to water vs sucrose.

**Figure 5:**
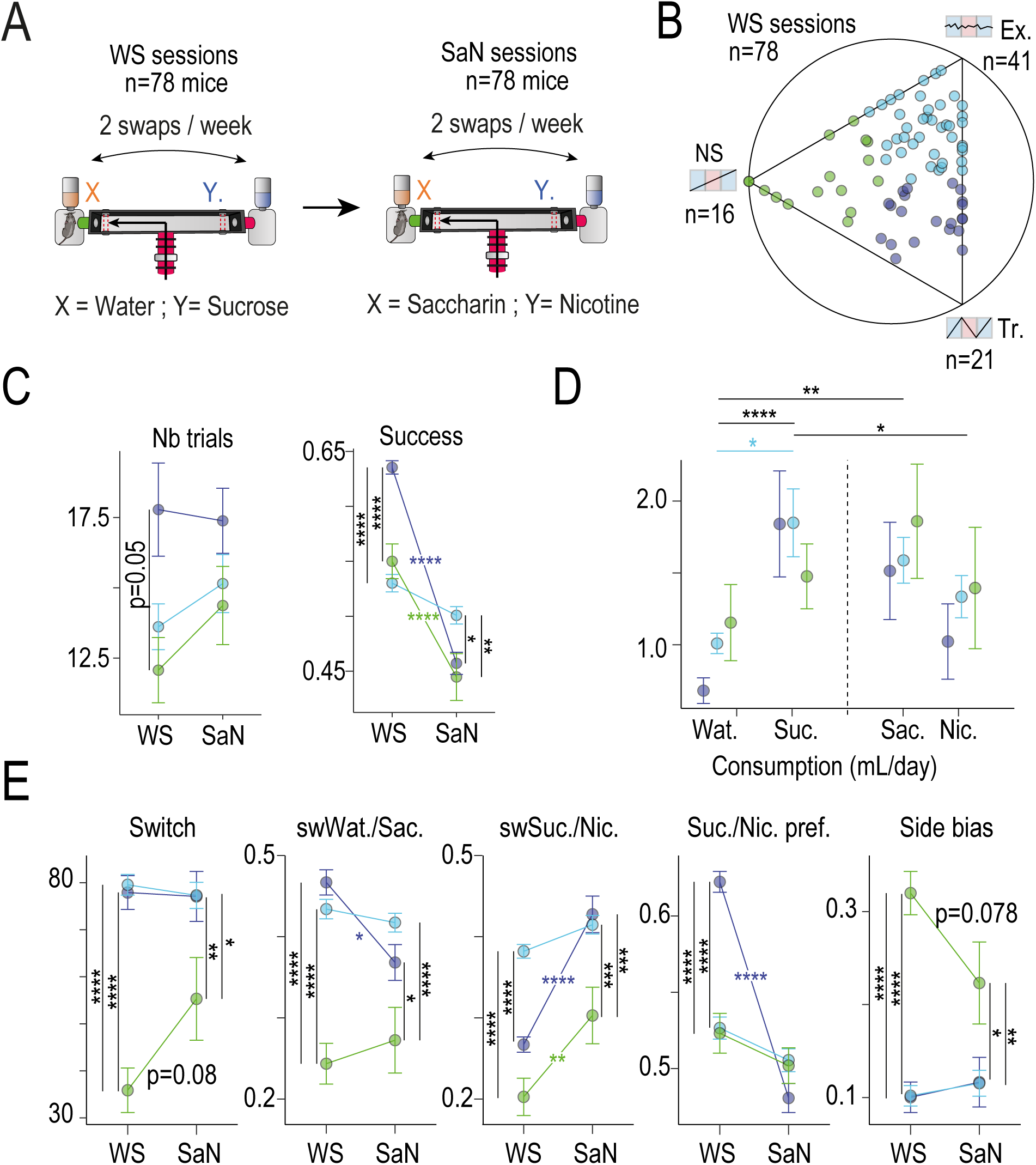
Predicting nicotine intake by sucrose-seeking strategy. **(A)** Experimental design **(B)** Position in the ternary plot of the 78 mice used in this analysis. These mice have been exposed to water-sucrose (WS) session and then to saccharine-nicotine (SaN) sessions. Ternary plot is obtained at the end of the WS sessions. **(C)** Mean number of T-maze entries per day (Nb trials) and mean Success for the three archetypes in WS and SaN periods (Wilcoxon tests with Holm correction, n indicated mice). **(D)** Mean consumption per animal and per archetype for water (Wat) or sucrose (Suc) and Saccharin (Sac) or Nicotine (Nic) sessions. **(E)** Variation in *Switch*, *SwWat*, *SwSuc, Pref*, *SideBias* (from left to Right) per archetypes and for WS and SaN periods.

#### Neural correlates of adaptability to the environment emerge after nicotine challenge

To determine how adaptation to environmental conditions influences the neural response to nicotine, we compared brain-wide cellular activation after an injection of nicotine or saline between mice living in Souris-City (at the end of SW session) and those living in standard home-cage housing (Figure 6A). Cellular activation to a challenge injection of saline or nicotine, measured as the numbers of cells expressing the immediate-early gene cFos, was compared between home-cage and Souris-City mice using the iDISCO+/ClearMap pipeline^43^. Mice living in Souris-City showed a greater differential activation across the brain in response to an acute nicotine challenge (Nicotine vs Saline injection, Figure 6B *Right*) than mice living under standard HC conditions (Figure 6B *Left*). Furthermore, mice in SC had overall greater brain-wide activation in response to an acute injection of nicotine than mice in HC (Nicotine HC vs. Nicotine SC, Figure 6C), including significant increases in cFos positive cells in regions associated with addiction, such as the medial prefrontal cortex, as well as in parts of the amygdala associated with anxiety and anxiety-like behavior. Indeed, voxel-by-voxel comparisons in these regions identify significant increases in activation only in the SC mice that received a nicotine injection (Figure 6D), suggesting that the adaptability required simply by living in a complex microsociety environment reorganizes brain-wide responsivity to nicotine. We next asked whether the distinct reward seeking profiles identified in SC, which are notably associated with distinct adaptations in the face of changing reward values, show differential activation of brain regions in response to nicotine injection. We correlated each mouse’s archetypal composition with their cFos-positive cell counts in response to saline or nicotine injection in 33 regions known to show alterations in cFos expression in response to nicotine ^44,45^. The number of cFos-positive cells did not correlate with archetypal composition following a saline injection in any of the brain regions studied (Figure 6E, *Right*). Following a nicotine injection, however, marked patterns of correlations are evident (Figure 6E *Left*): mice that show adaptations in response to ΔV in the T maze, namely the Tracker and the Non-Switcher archetypes, showed strong positive correlations across one third of the pre-defined regions, whereas the Explorer mice showed largely negative correlations. Increases in cellular reactivity to this nicotine challenge may thus be linked to behavioral and choice adaptability in the face of changing reward values.

**Figure 6:**
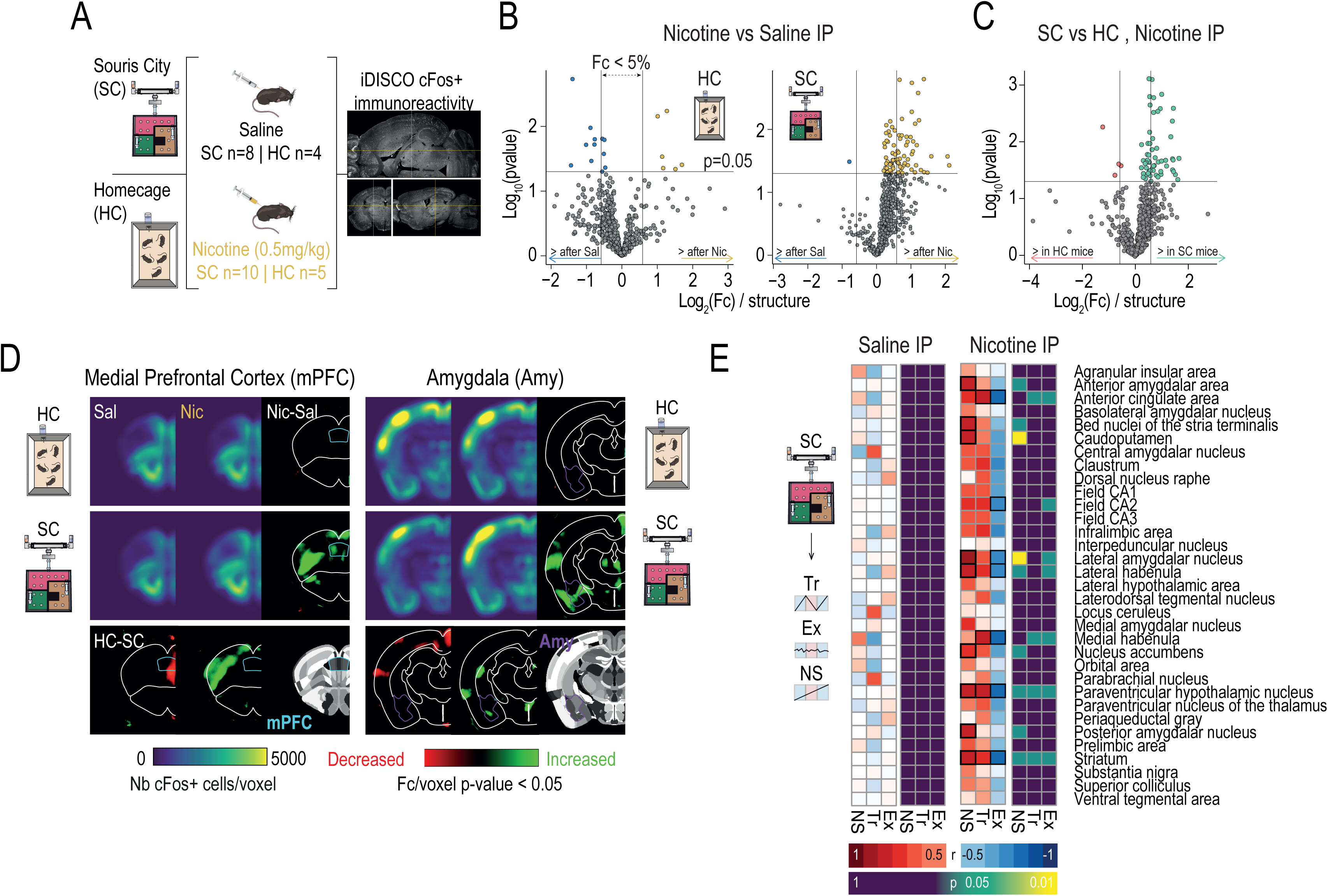
Differential nicotine response as a function of the environment and decision-making strategy. **(A)** Brain-wide activity mapping after saline or nicotine injection in mice living in standard home cages (HC) or Souris-City (SC) revealed by iDISCO brain clearing and Clearmap activity. **(B)** Mice in HC conditions do not show significant activation differences between saline and nicotine injections, while mice in SC show a shift toward increased cFos expression following a nicotine injection when compared to a saline one. **(C)** Comparison between nicotine induced cFos expression in SC and HC mice reveals that SC mice show greater numbers of activated cells in response to nicotine than mice raised under standard conditions. **(D**) Grouped heatmaps show average cFos expression density in the PFC (*Left*) and the amygdala (*Right*). P-value maps highlight areas where significant between-groups differences can be appreciated. **(E)** Correlations between cFos expression and distance to the archetype in saline and nicotine-injected SC animals *Left*: minimal differences in cFos expression are observed between mouse profiles in SC in response to a saline injection. *Right*: patterns of expression across brain regions associated with cognitive and reward functions that become apparent after challenging the dopaminergic system in mouse profiles in SC.

We observed altered reactivity to nicotine across many brain regions, notably including dopamine receptive regions (e.g. the striatum), and regions that regulate dopamine neuron firing (e.g. the lateral habenula). Thus, we next asked if adaptation to the SC environment would also lead to alterations in VTA DA neuron activity. We first used *in vivo* juxtacellular recording to assess the spontaneous activity of VTA DA neurons between mice immediately upon their exit from SC and compared this with HC mice. Because the exposure to sucrose in the SC environment could, in and of itself, alter DA activity, two groups of HC mice were studied: one with access only to water (HC/Wat), and one with access to a 5% sucrose drinking solution (HC/Suc) (Figure 7A). Both the firing rate and the bursting activity (%SWB) of VTA DA cells were significantly higher in mice with access to sucrose as compared to water in HC condition (Figure 7B). In Souris-City, VTA DA cells showed intermediate firing rate and an overall reduced bursting rate compared to the two HC conditions. These results confirm that while sucrose exposure alters DA neuron activity, it is not sufficient to explain the differences we find in SC. Thus, both sucrose exposure and the environment interact to modulate VTA DA cells activity^7^.

**Figure 7:**
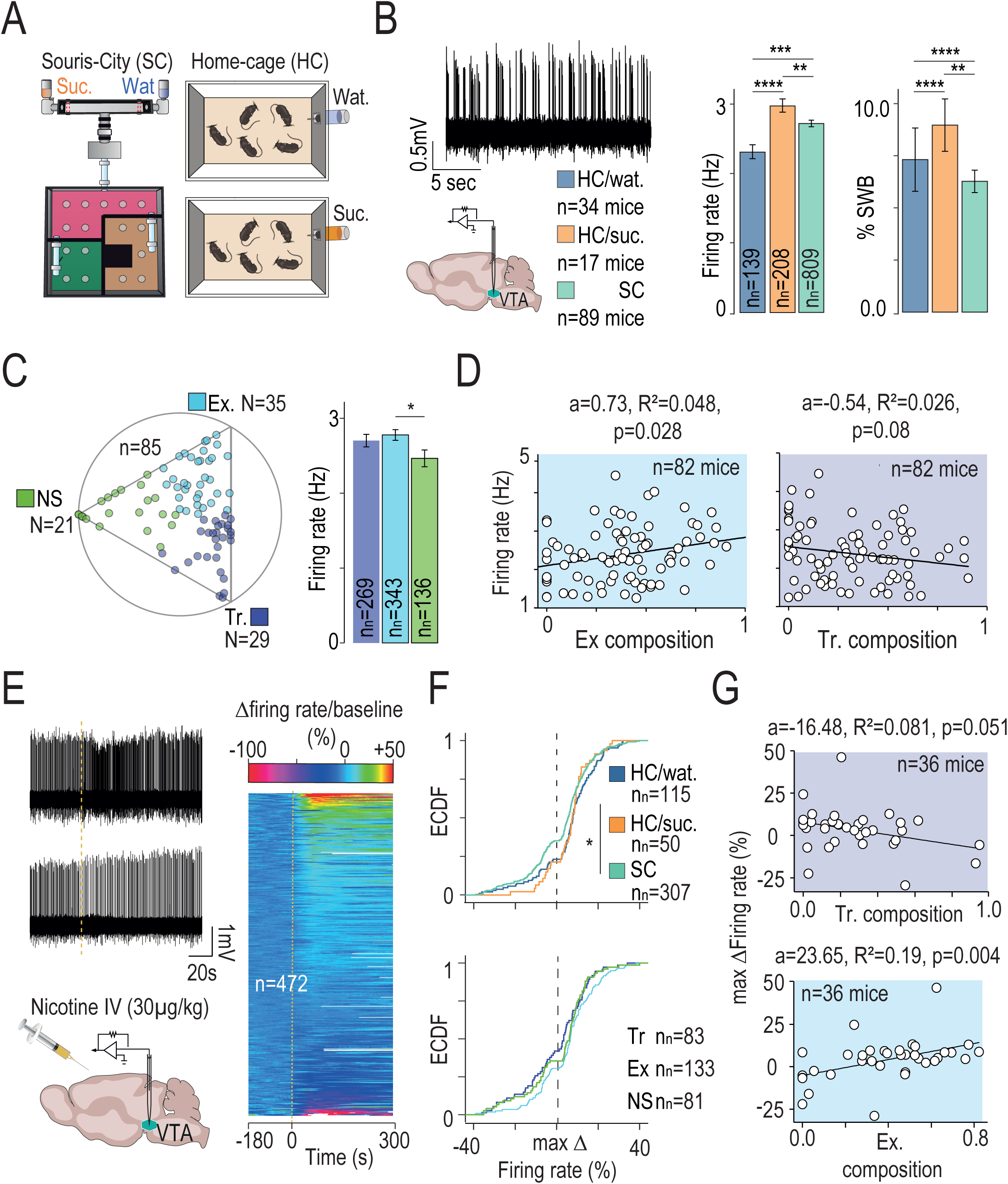
Dopamine neuron firing is modulated by both the environment and mouse personality. **(A)** Spontaneous activity of VTA dopaminergic (DA) neurons recorded in mice living in standard home cage (HC) with access either to water (Wat) or to a 5% sucrose drinking solution (Suc) or in Souris-City (SC). **(B)** Mean firing rate are different in the three groups (Kruskal-Wallis (df=2, p=0.017) and post hoc Wilcoxon tests with Holm correction, *p<0.05, n indicated number of mice, nn indicated number of neurons). **(C)** *Left:* Position in the ternary plot of the n=85 mice used in this analysis. *Right:* Firing rate in Hz according to the archetype (Wilcoxon tests with Holm correction, n=748 neurons). **(D)** Correlation (linear regression, a indicated the slope estimate, R^2^ the Adjusted R-squared and p the p-value) between Explorer (Ex, *Left*) or Tracker (Tr, *Right*) composition and median firing rate per mice (n= 82 mice, with a minimum of 3 cells per mice). **(E)** *Left*: Intravenous (i.v.) injections of nicotine (Nic; 30 µg/kg) induce activation (upper panel) or inhibition (lower panel) of distinct VTA DA neurons in anesthetized mice (representative recordings). *Right*: Responses of VTA DA neurons after nicotine injection. Responses are rank-ordered based on the response to nicotine, from the most excited to the most inhibited (top to bottom of the graph). Color scale indicates variation in firing rate amplitude. **(F)** Empirical cumulative distribution of the response to nicotine (variation in firing rate). (*Top*) Neurons recorded in mice living in Souris-city, in home cage (HC) with water (HC/Wat) or with sucrose drinking solution (HC/Suc) (Kruskal-Wallis (df=2, p=0.011) and post hoc Wilcoxon tests with Holm correction, *p<0.05). (*Bottom*) Neurons recorded in SC mice according to their Tr, Ex and NS respective profiles (Kruskal-Wallis (df=2, p=0.059)). **(G)** Correlation between Tracker (Tr, *Top*) or Explorer (Ex, *Bottom*) composition and median response in firing rate per mice (n=36 mice, with a minimum of 5 cells per mice).

To dissect whether the differences that we observed in spontaneous VTA DA neuron activity in SC mice are related to mouse personalities, we categorized the SC mice according to their archetypal profiles (Figure 7C, *Left*, n = 85 mice). Comparing basal DA neuron activity between the archetype groups revealed a significant difference in firing rate between the Explorers and Non-Switchers (Figure 7C, *Right*). Averaging dopaminergic activity per mouse revealed a correlation between the proximity to the archetype and the firing rate for Explorers (Figure 5D, *Right*, p=0.028) but not for Trackers (Figure, 7D, *Left*, p=0.08) or Non-Switchers (not shown, p = 0.8). We then measured nicotine-evoked responses in anesthetized animals to investigate whether the environmental condition impacts the sensitivity of DA neurons to nicotine (Figure 7E). VTA neurons are not a uniform population and we have shown that nicotine concomitantly activates and inhibits two distinct subpopulations of DA neurons^46,47^. Therefore, when recording the response of a population of DA neurons to nicotine, it can be organized from the most inhibited to the most excited (Figure 7E, *Right*). Comparison of the distributions according to the environments reveals differences (Figure 7F, *Top,*HC/suc versus SC p=0.013) while comparison of the distributions according to the profiles observed in SC (Figure 7F, *Bottom*) did not highlight reliable differences (p=0.059). On the other hand, we observed clear correlations, between the proximity to the Tracker and Explorer archetypes and the median response per animal (Figure 7G).

Together, these data indicate the existence of correlations between sucrose-seeking patterns in a choice task and the response to nicotine, suggesting that adaptations in dopaminergic circuit function emerge between mice different personalities identified by their decision-making strategies.

## Conclusion

In this study, we found that a complex social environment drives individuation of mouse reward-seeking strategy, even when tested isolated from their peers. These differences are further linked to variations in behavioral traits, or “personality”, in the activity of the dopaminergic system, and in the response to nicotine. By using a reinforcement learning (RL) model trained on a specific choice situation (water vs. sucrose) we were able to accurately predict the animals’ behavior in a different situation involving a choice between saccharine alone and saccharine with nicotine. These results suggest that in a complex social environment (i) animals adopt distinct foraging strategies, (ii) these strategies reflect individual personality traits and the state of neural circuits, and (iii) an individual’s strategy can be indicative of their response to addictive substances. Overall, the study highlights how harnessing inter-individual variability in behavior and its underlying mechanisms, particularly in the context of addiction research, can unravel more complex and nuanced relationships between neural circuits and behavior than would be possible by assuming that all mice should respond uniformly to a task. The use of large naturalistic environments with automated data capture provides a valuable tool for studying individual variations and their implications for the development and maintenance of personality profiles and susceptibility to substances of abuse like nicotine.

There is ongoing debate regarding the emergence of inter-individual variation and the concept of personality. Our experiments demonstrate that when animals are placed in controlled, yet naturalistic environments, distinct profiles, marked by clusters of behavioral traits, emerge and remain stable over extended periods, which can be considered the definition of personality. Being able to test animals in isolation in a simple choice test eliminates direct social influences and mitigates issues related to multicollinearity and co-dependence in the main environment. This approach also reveals the complexity of the underlying mechanisms. The broad concept of cerebral plasticity and the embedding of an individual in an environment allow for the consideration of each individual’s singularity. This singularity is manifested through differences in neural connections and activities, but that alone is insufficient to explain inter-individual variability and the emergence of stable differences. Connections and activities themselves are modified by an individual’s history and by the environment in which they live ^48^. No two animals can live in exactly the same environment, as how each individual engages with their physical surroundings and social milieu will be different as a function of their bodily constraints and personality traits (e.g. their level of sociability). Thus, when animals live in the same physical space they necessarily adapt differently. Studies involving grouped animals have demonstrated this adaptability, showing that initial diversity reemerges when animals with similar profiles are grouped together^7^. The asymmetry of the environment, which includes social relations and competition for resources, facilitates the development of individuality, with both shared and non-shared components within each individual’s perceived environment^4,49^. When an animal is exposed to a test, isolated from the others, the differences in strategies that can be seen in the test reflect the differences that exist in the main environment and are the consequences of each individual’s adaptations to both that environment and its co-living congeners. In our study, this is reflected in differences in dopaminergic activity, responses to nicotine, and strategies employed in the T-maze.

Differences in the T-maze can be captured by reinforcement learning models. Variances observed in a specific choice context (water-sucrose) can be explained by discrepancies in learning rates and exploration parameters. These models accurately predict how responses evolve when the difference in value between proposed rewards is changed. These results confirm the validity of these computational approaches to account for behavior and the use of latent variables to describe such complex phenomena. Notably, the behavior of non-switchers animals becomes particularly intriguing within this context. While an initial judgment based on their side bias could conclude that these mice are insensitive to differences in value (i.e. low β), our modeling findings indicate that these animals in fact have a *high* exploitation level (high β), albeit paired with a very low learning rate, meaning that they update their representations slowly. When looking closer at their behavior, we indeed find that their bias is for the side where they encounter sucrose early on. This gradual and slow adaptation of value, combined with a strong bias toward the highest value and an environment characterized by rapid position changes of significant rewards, allows for the emergence of this seemingly “maladaptive” behavior. In a previous study, we were able to demonstrate that modifying the social environment, i.e regrouping together non-switching animals, resulted in a fast re-adaptation of both the strategy and the dopamine firing pattern, suggesting an important impact of adaptation to the local environment ^7,20^. If all latent variable values are a consequence of adaptation to the environment (both fixed and social), the remaining question pertains to the constraint that an individual must adapt in order to reduce its learning rate.

Finally, the differentiation of profiles observed in our study is closely linked to the unique patterns of spontaneous activity and response of dopaminergic neurons to nicotine. To fully understand the complex relationship between behavior and neural network activity, and to gain insight into drug susceptibility, it is imperative to systematically categorize individual differences. While it may be difficult to identify a single circuit responsible for behavior using current approaches, we can propose that the dopaminergic system, among others, occupies a central position at the intersection of several factors. On the one hand, the activity of the dopaminergic system is significantly influenced by an individual’s personal history, social interactions, and cumulative experiences^7,20^. These factors shape the functional state of the dopaminergic system and ultimately influence its involvement in various behavioral processes. On the other hand, the dopaminergic system itself plays a central role in defining and regulating specific parameters related to decision making and behavioral expression^22,23,50,51^. It acts as a modulator, influencing the salience and motivational value assigned to stimuli and guiding the selection of appropriate actions in response to environmental cues. This central position of the dopaminergic system, shared by numerous brain regions, allows us to conceptualize a neural basis for the notion of individuality^20^. Thus, in this sense, our results suggest that a dynamic feedback loop exists between environmental encounters and the dopaminergic system which continually defines animal traits and adaptability to future challenges.

By recognizing the complex interdependencies between neural activity, behavior, and environmental influences, we can gain a deeper understanding of the multifaceted nature of individual differences. This understanding is critical not only for unraveling the mechanisms underlying behavior, but also for elucidating the factors that contribute to susceptibility to drugs and addictive substances. Ultimately, a comprehensive exploration of these interrelated factors will allow us to develop more targeted and personalized approaches to addiction research and behavioral intervention.

## Acknowledgements

This work was supported by the Centre National de la Recherche Scientifique CNRS UMR 8246 and 8249, INSERM U1130, the Foundation for Medical Research (FRM, Equipe FRM DEQ2013326488 to P.F), the French National Cancer Institute Grant TABAC-16-01, TABAC-19-020 and SPA-21-002 (to P.F.), French state funds managed by the ANR (ANR-19-CE16-0028 Bavar to PF and NR). LMR was supported by a NIDA–Inserm Postdoctoral Drug Abuse Research Fellowship. Fourth-year PhD fellowship from Fondation pour la Recherche Médicale (FDT201904008060 to SM). Fourth-year PhD fellowship from the Biopsy Labex (CN). We are grateful to the animal facilities (IBPS) and Otilia de Oliveira, Noemie Karakaplan-Dherbe and Emilie Tubeuf at ESPCI animal facilities.

## Author contributions

Conceptualization, SLF, LMR and PF; Methodology, SLF, LMR, NT, FM, NR and PF; Validation: SLF, LMR, FM, NR, AM and PF; Formal Analysis, SLF, LMR, RJ, SD, ND, YL and PF; Investigation, SLF, NT, BH, LMR, ST, SM, CN, CV, LR, AMC, TT, FM, Original Draft SLF, LMR, CV, FM, AM and PF; Review & Editing all authors, supervision PF, AM, FM, NR, Funding acquisition PF

## Methods

### Animals

Experiments were performed on adult C57Bl/6Rj Wild-Type (Janvier Labs, France) mice. Male mice, from 8 to 10 weeks old at the beginning of the experiment, weighing 25-35 grams, were used for all the experiments. They were kept in an animal facility where temperature (20 ± 2°C) and humidity were automatically monitored and a circadian light cycle of 12/12-h light-dark cycle was maintained. All experiments were performed in accordance with the recommendations for animal experiments issued by the European Commission directives 219/1990, 220/1990 and 2010/63, and approved by Ethic Committee. All mice were implanted under anesthesia (isoflurane 3% – Iso-Vet, Piramal, UK), with an RFID chip subcutaneously inserted in the back.

### Souris City setup

#### Setup

Souris City combines a large environment (the social cage 1*1 m) where groups of male mice live for extended periods of time in semi-natural conditions, and a test-zone where mice have a controlled access to specific areas for drinking. Souris City was house-designed and built by TSE Systems (Germany). Mice were tagged with RFID chips, allowing automatic detection and controlled access to the different areas. Animals were living under a 12h/12h dark-light cycle (lights on at 7am) and had access to food ad libitum. The social cage is divided into four sub-compartments. These different sub-compartments are equipped with RFID antennae on the floor and are connected through tubes that are also equipped with antennae. Therefore, each transition from one sub-compartment to another was associated with a detection of the animal by the two antennae of the transition tube ^7^.

The social cage is connected to the test zone by a gate, which is a key element of the setup. The gate (TSE Systems, Germany) is composed of three doors with independent automatic control. It allows for the selection of animals and grants control over their access to the test zone. In the test zone, mice are isolated from their peers and conduct the test on their own, without any intervention from the experimenter. The test consists in a T-maze choice task ^52^. Since the T-maze was the only source of water, animals were motivated to perform the test. The T-maze gives access to two home-cages, one on each side (left and right), with a drinking bottle in each. The bottles contained either water, sucrose, saccharin or nicotine. The system was configured in such a way that animals performed a dynamic foraging task. The reward value of the bottle content is changed every 3-4 days to evaluate whether mice were able to track the highest reward. Automating the task in this manner minimizes the need for human involvement, thereby mitigating limitations such as cost and time constraints associated with human assessment. Moreover, it eliminates the potential risks of inducing stress or disturbing the natural cycle of the animals.^53–56^. Simple rules were used to automatize the test. When a mouse accesses one liquid bottle, an infra red-light beam is interrupted in that particular arm, resulting in the closing of the access to bottle on the opposite side. A Plexiglas cylinder descends, preventing access to the bottle. To initiate a new trial, mice must exit the T-maze, which triggers the reopening of the feeders. For the analysis in T-maze, throughout the paper, only mice accessing the T-maze more than 5 times per day are included in the analyses (i.e. 281/295). The only exception is in Figure 7B for the analysis of dopamine neuron activity after exposure to SC, where all mice are considered (in this group, 4 out of 89 mice made no more than 5 trials per day).

#### In vivo juxtacellular recordings of VTA DA neurons

Mice were deeply anaesthetized with chloral hydrate (8%), 400 mg/kg IP, supplemented as required to maintain optimal anesthesia throughout the experiment. The scalp was opened and a hole was drilled in the skull above the location of the VTA. Extracellular recording electrodes were constructed from 1.5 mm O.D. / 1.17 mm I.D. borosilicate glass tubings (Harvard Apparatus) using a vertical electrode puller (Narishige). Under microscopic control, the tip was broken to obtain a diameter of approximately 1 µm. The electrodes were filled with a 0.5% NaCl solution containing 1.5% of Neurobiotin tracer (AbCys) yielding impedances of 6-9 MΩ. Electrical signals were amplified by a high-impedance amplifier (Axon Instruments) and monitored through an audio monitor (A.M. Systems Inc.). The signal was digitized, sampled at 25 kHz and recorded using Spike2 software (Cambridge Electronic Design) for later analysis. The electrophysiological activity was sampled in the central region of the VTA (coordinates from bregma: 3.1 to 4 mm AP, 0.3 to 0.7 mm ML, and 4 to 4.8 mm DV from the brain surface). Individual electrode tracks were separated from one another by at least 0.1 mm in the horizontal plane. Spontaneously active DA neurons were identified based on previously established electrophysiological criteria ^25,57^. Intravenous administration of saline (H_2_O with 0.9% NaCl) or nicotine at a dose of 30 µg/kg (4.16 mg/kg, free base) was carried out through a catheter (30G needle connected to polyethylene tubing PE10) connected to a Hamilton syringe, into the saphenous vein of the animal.

#### Solution in T-maze experiment

In the T-maze, mice were presented with i) two bottle of water ii) one bottle of water and one bottle of sucrose (5%, Sigma Aldrich) iii) one bottle of saccharine solution (2%, Sigma Aldrich) and one bottle of nicotine (100 µg/ml free base, Sigma Aldrich) plus saccharine (2%) solution diluted in water (adjusted to pH ∼7.2 with NaOH).

#### Brain clearing and activity mapping

Expression of the immediate early gene *c-fos* is used as a marker for cellular activation, since it is rapidly induced following a stimulus, readily immunolabeled in optically cleared brains, and spatially restricted to the cell nucleus giving high signal-to-noise for automated counting.

#### Experimental design and perfusion

Mice were injected with i.p. saline or nicotine and kept in a dim, quiet room for one hour before perfusion to minimize off-target cFos expression. Mice were then perfused with 1X PBS followed by 20mL of 4% paraformaldehyde (PFA, Electron Microscopy Services). Brains were carefully dissected from the skull, and stored in PFA overnight. Brains were stored in PBS with 0.01% Sodium Azide (Sigma-Aldrich, Germany) until clearing.

##### iDISCO+ whole brain immunolabeling

Whole brain clearing and immunostaining was performed following the iDISCO+ protocol previously described previously ^58^ with minimal modifications. All the steps of the protocol were done at room temperature with gentle shaking unless otherwise specified. All the buffers were supplemented with 0.01% Sodium Azide (Sigma-Aldrich, Germany) to prevent bacterial and fungal growth. Briefly, perfused brains were dehydrated in an increasing series of methanol (Sigma-Aldrich, France) dilutions in water (washes of 1 hour in methanol 20%, 40%, 60%, 80% and 100%). An additional wash of 2 hours in methanol 100% was done to remove residual water. Once dehydrated, samples were incubated overnight in a solution containing a 66% dichloromethane (Sigma-Aldrich, Germany) in methanol, and then washed twice in methanol 100% (4 hours each wash). Samples were then bleached overnight at 4 C in methanol containing a 5% of hydrogen peroxide (Sigma-Aldrich). Rehydration was done by incubating the samples in methanol 60%, 40% and 20% (1 hour each wash). After methanol pretreatment, samples were washed in PBS twice 15 minutes and 1 hour in PBS containing a 0.2% of Triton X-100 (Sigma-Aldrich) and further permeabilized by a 24 hours incubation at 37C in Permeabilization Solution, composed by 20% dimethyl sulfoxide (Sigma-Aldrich), 2.3% Glycine (Sigma-Aldrich, USA) in PBS-T. In order to start the immunostaining, samples were first blocked with 0.2% gelatin (Sigma-Aldrich) in PBS-T for 24 hours at 37C, the same blocking buffer was used to prepare antibody solutions. Brains were incubated with anti c-Fos primary antibody (Synaptic systems 226-003) for 10 days at 37°C with gentle shaking, then washed in PBS-T (twice 1 hour and then overnight), and finally newly incubated for 10 days with secondary antibodies. Secondary antibodies raised in donkeys, conjugated to Alexa 647 were used (Life Technologies). After immunostaining, the samples were washed in PBS-T (twice 1 hour and then overnight), dehydrated in a methanol/water increasing concentration series (20%, 40%, 60%, 80%, 100% one hour each and then methanol 100% overnight), followed by a wash in 66% dichloromethane – 33% methanol for 3 hours. Methanol was washed out with two final washes in dichloromethane 100% (15 min each) and finally the samples were cleared and stored in dibenzyl ether (Sigma-Aldrich) until light sheet imaging.

##### Light sheet microscopy

The acquisitions were done on a LaVision Ultramicroscope II equipped with infinity-corrected objectives. The microscope was installed on an active vibration filtration device, itself put on a marble compressed-air table. Imaging was done with the following filters: 595/40 for Alexa Fluor-555, and -680/30 for Alexa Fluor-647. The microscope was equipped with the following laser lines: OBIS-561nm 100mW, OBIS-639nm 70mW, and used the 2nd generation LaVision beam combiner. The images were acquired with an Andor CMOS sNEO camera. Main acquisitions were done with the LVMI-Fluor 4X/O.3 WD6 LaVision Biotec objective. The microscope was connected to a computer equipped with SSD drives to speed up the acquisition. The brain was positioned in sagittal orientation, cortex side facing the light sheet, to maximize image quality and consistency. A field of view of 1000 x 1300 pixels was cropped at the center of the camera sensor. The light sheet numerical aperture was set to NA-0.03. The 3 light sheets facing the cortex were used, while the other side illumination was deactivated to improve the axial resolution. Beam width was set to the maximum. Laser powers were set to 40-60% (639nm). The center of the light sheet in x was carefully calibrated to the center of the field. z steps were set to 6mm. Tile overlaps were set to 10%. The whole acquisition took about 1h per hemisphere. At the end of the acquisition, the objective was changed to a MI PLAN 1.1X/0.1 for the reference scan at 488nm excitation (tissue autofluorescence). The field of view was cropped to the size of the brain, and the z-steps are set to 6mm, and light sheet numerical aperture to 0.03 NA. It was important to crop the field of view to the size of the brain for subsequent alignment steps.

##### Computing Resources

The data were automatically transferred every day from the acquisition computer to a Lustre server for storage. The processing with ClearMap was done on local workstations, either Dell Precision T7920 or HP Z840. Each workstation was equipped with 2 Intel Xeon Gold 6128 3.4G 6C/12T CPUs, 512Gb of 2666MHz DDR4 RAM, 4×1Tb NVMe Class 40 Solid State Drives in a RAID0 array (plus a separate system disk), and an NVIDIA Quadro P6000, 24Gb VRAM video card. The workstations were operated by Linux Ubuntu 20.04LTS. ClearMap 2.0 was used on Anaconda Python 3.7 environment.

##### ClearMap Fos+ cell counting

Tiled acquisitions of Fos-immunolabeled iDISCO+ cleared brains scanned with the light sheet microscope were processed with ClearMap 2^43^ to generate both voxel maps of c-Fos cell densities, as well as region-based statistics of cell counts. Stitched images were processed for background removal, on which local maxima were detected to place initial seeds for the cells. A watershed was done on each seed to estimate the volume of the cell, and the cells were filtered according to their volume to exclude smaller artefactual maxima. The alignment of the brain to the Allen Brain Atlas was based on the acquired autofluorescence image using Elastix ^59^ (https://elastix.lumc.nl). Filtered cell’s coordinates were transformed to their reference coordinate in the Allen Brain Atlas common coordinate system ^60^. For voxel maps, spheres of 375mm diameter were drawn on each filtered cell. P-value maps of significant differences between groups were generated using Mann-Whitney U test (SciPy implementation). Aligned voxelized datasets from each group of animals were manually inspected to identify the regional overlaps of p-value clusters, and volcano-plots of regional counts were generated.

#### Decision Model

Data choice (i.e Left or Right) from all mice in the T-maze were modelled and fitted with a standard RL model ^41^. The model uses the sequence of choices and outcomes (the reward) to estimate the expected value of each option for every trial. After each trial, the value associated with the chosen option was updated according to the classical delta rule: V_R_(t+1)=V_R_(t)+αδ(t) and δ(t)=R_R_(t)-V_R_(t) where δ(t) is the reward prediction error (RPE), the difference between the expected value and the received reward, i.e the reward prediction error (RPE). V_i_={V_L_,V_R_} is set to 1 when reward is water and 3 when reward is sucrose. For modeling, the expected values are set to zero at the beginning of the experiment. The learning rate α determines how rapidly the estimate of expected value is updated. Given expected values V_i_ for both options, the probability of choosing the right option P_R_(t) is computed using a SoftMax rule defined by P_R_(t) =1/(1+exp(-[ß(V_R_(t)-V_L_(t)) + χ(C_R_(t-1)-C_L_(t-1))]) with two parameters β and χ. The inverse temperature parameter β represents the sensitivity to the difference of values V_i_={V_L_,V_R_}, it reflects how much the difference in total value between the two options (ΔV) translates into more or less preference for the best option in a given gamble. With a small β, choices have low sensitivity for ΔV, with the extreme case of a null β where both options have the same probability to be selected. On the contrary, a large β indicates a high sensitivity to ΔV, with an infinite beta indicating that options associated with higher reward probabilities are always selected. The choice perseveration parameter χ captures short-term tendencies (previous choice) to perseverate or alternate (when positive or negative, respectively). This tendency to alternate is independent of the reward history ^42^, and thus does not depend on ΔV. The free parameters of the model (α,β,χ) were fitted by maximizing the data likelihood. Given a sequence of choice c = c_1…T_, data likelihood is the product of their probability given by the SoftMax choice rule ^61^. We used the optim function in R to perform the fits, with the constraints that α ∈]0,10], β ∈]0,5] and χ ∈]0,5].

#### Statistical analysis

All statistical analyses were computed using R (The R Project, version 4.0.0) and Python with custom programs. Results were plotted as a mean ± s.e.m. The total number (n) of observations in each group and the statistics used are indicated in figure legends. Classical comparisons between means were performed using parametric tests (Student’s T-test, or ANOVA for comparing more than two groups when parameters followed a normal distribution (Shapiro test P > 0.05), and non-parametric tests (here, Wilcoxon or Mann-Whitney) when the distribution was skewed. Multiple comparisons were corrected using a sequentially rejective multiple test procedure (Holm). Probability distributions were compared using the Kolmogorov–Smirnov (KS) test, and proportions were evaluated using a chi-squared test (χ²). All statistical tests were two-sided except for the nicotine exposure session experiment (Figure 5) where statistical tests were one-sided (we test hypotheses driven by nicotine effect and model). P > 0.05 was considered not to be statistically significant. For archetypal analysis, all computations and graphics have been done using the statistical software R and the archetype package (version 2.2-0.1). Briefly, the archetypal analysis finds the matrix Z of k m-dimensional archetypes (k is the number of archetypes), given an n × m matrix representing a multivariate data set with n observations (n = number of animals) and m attributes (here m = 5, consisting of the level of global switching (*Switch*), the variables *SwWat* and *SwSuc* that represent the probability of switching sides if the previous choice was water or sucrose, respectively, the *Pref* and *SideBias* that assess preference and side bias by comparing the choices (% of sucrose choice) between each session. Z is obtained by minimizing || X-α Z^T^ ||_2_, with α the coefficients of the archetypes (α_i,1..k_ ≥0 and ∑α_i,1..k_ = 1), and ||.||_2_ a matrix norm. The archetype is also a convex combination of the data points Z=X^T^δ with 8≥ δ and their sum must be 1 ^62^. The α-coefficient depicts the relative archetypal composition of a given observation. For k = 3 archetypes and an observation i, α_i,1_ ; α_i,2_ ; α_i,3_ ≥ 0 and α_,1_ +α_i,2_ + α_i,3_ = 1. A ternary plot can then be used to visualize data. (α_i,1_ ; α_i,2_ ; α_i,2_) are used to assign individual behavior to its nearest archetype (i.e, k max(α_i,1_ ; α_i,2_ ; α_i,3_)). α_i,j_ are also used as variable to estimate population archetypal composition. Archetypal composition correspond to α (0 ≤ α_i,j_ ≤ 1). Pure archetype corresponds to 1, the archetypal composition decreases linearly with increasing distance from the archetype, 0 correspond to points on the opposite side.

#### Statistics and Reproducibility

All experiments were replicated with success.

#### Data Availability

All the data that support the findings of this study can be found in the Source Data file provided with the paper. If necessary, the raw data from the online behavioral experiment (i.e the trajectories) are available from the corresponding author.

#### Code Availability

All codes used to run the analysis are available from the authors upon request.

---

**Supp Figure 3:**
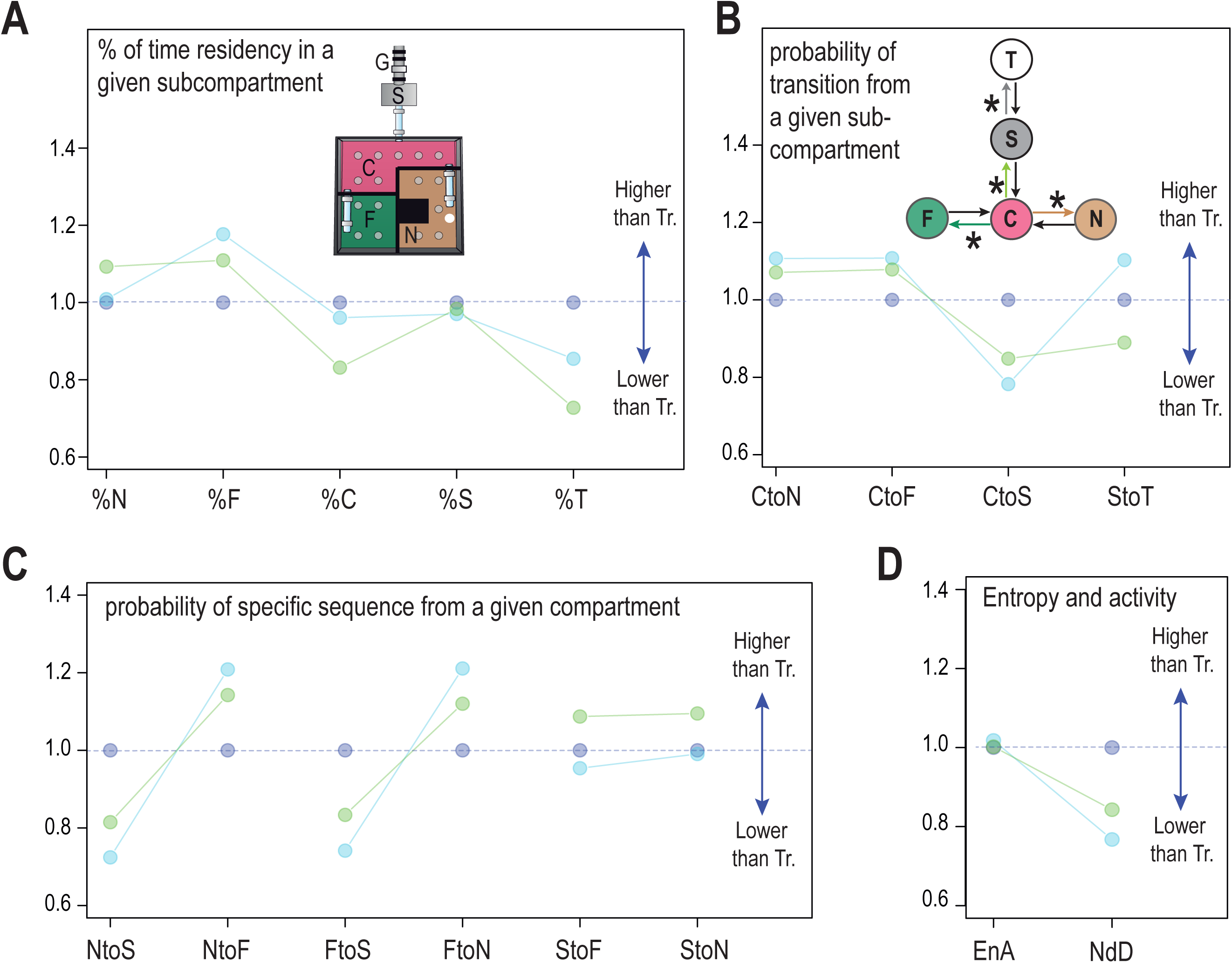
Comparison of the main environment variables depending on the archetype: Data are normalized to mean Tracker value. (A) variation in Occupancy, i.e from left to right percent time in Nest compartment (%N), percent time in Food compartment (%F), percent time in Center compartment (%C), percent time in Stair compartment (%S), percent time in T-Maze compartment (%T). (B) First order probability transitions. *Inset:* Transition of interest are labeled with a star. Center to Nest (CtoN), Center to Food (CtoF), Center to Stair (CtoS), Stair to T-maze (StoT). (C) Second order transition (probability) : Nest to Stair (NtoS), Nest to Food (NtoF), Food to Stair (FtoS), Food to Nest (FtoN), Stair to Food (StoF), Stair to Nest (StoN). (D) Activity Levels : Entropy (EnA), Number of Detections (NbD).

**Supp Figure 4:**
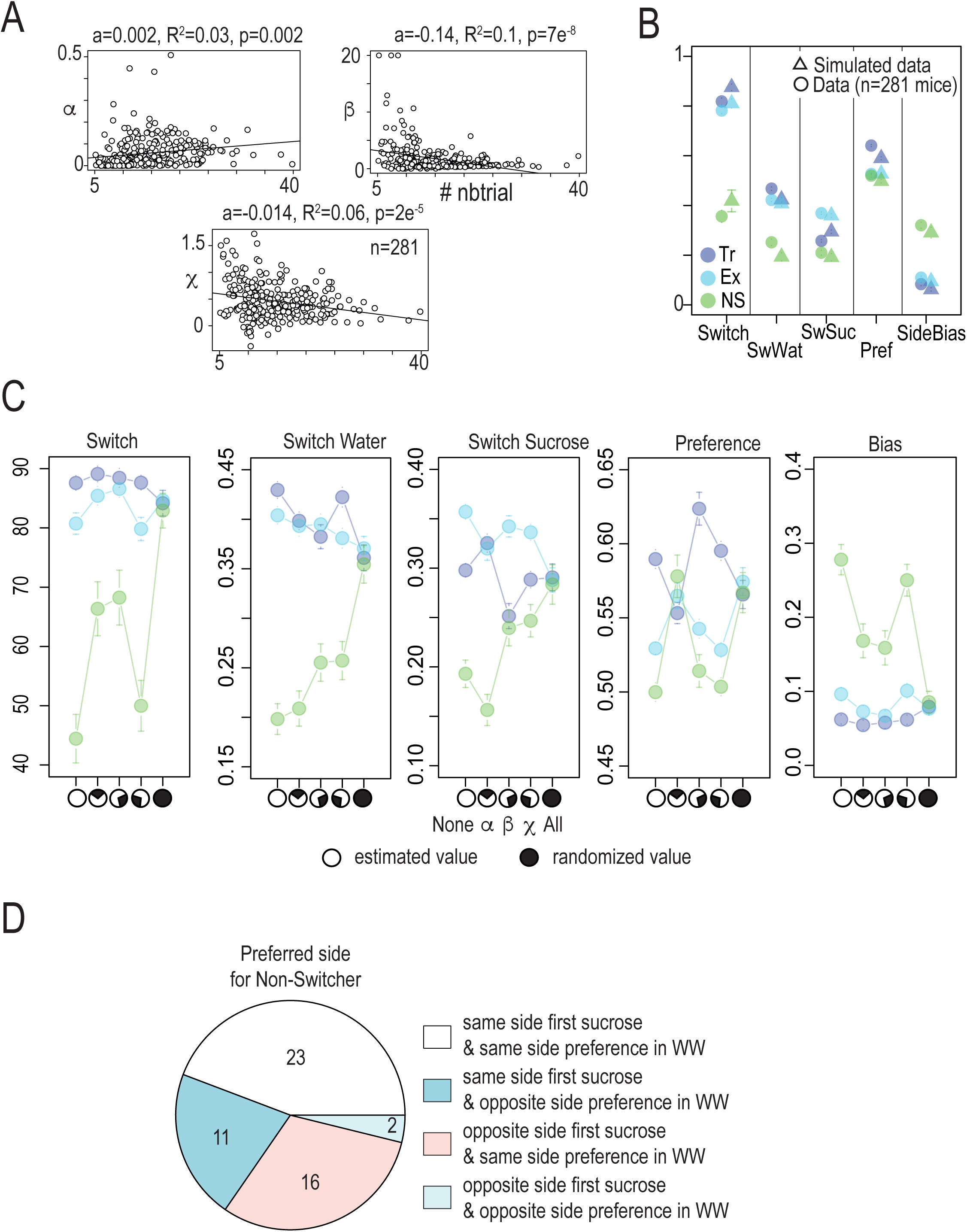
**(A)** Correlation (linear regression, a indicated the slope estimate, R^2^ the Adjusted R-squared and p the p-value) between the mean number of T-maze entry per day (# nbtrial) and χ (*Top Left*), β (*Bottom*) and α (*Top right*) (n=281 mice). **(B)** Comparison of the mean of five variable (*Switch*, *SwWat*, *SwSuc, Pref*, *SideBias*) for Tr., Ex. and NS archetype obtained for data (o) and for a model sequence (Δ) of 300 choices (6 sessions of 50 choices) simulated with fitted values of α, β and χ. **(C)** Attribution study in which each of the latent variables is manipulated independently from the other to assess its contribution to the *Switch*, *SwWat*, *SwSuc, Pref*, *SideBias* variables, for the three archetypes. The legend symbols represent different simulation conditions. Empty circle: Each mouse is simulated using its estimated values of latent variables α, β and χ. Black circle: Each mouse is simulated with a random selection of latent variables. α, β and χ. White and Black Circle: Each mouse is simulated with two of its estimated values and one is chosen randomly from the corresponding latent variables, respectively α, β or χ. **(D)** Percentage of Non-Switcher mice (n=52) with a preferred side in WS session that correspond to : (white) same first sucrose choice side in WS session and same side preference in WW session, (blue) same first sucrose choice side in WS session and opposite side preference in WW session, (pink) opposite first sucrose choice side in WS session and same side preference in WW session and (light blue) opposite first sucrose choice side in WS session and opposite side preference in WW session.

## Bibliography

1. Deroche-Gamonet, V., Belin, D. & Piazza, P. V. Evidence for addiction-like behavior in the rat. Science (New York, N.Y.) 305, 1014–1017 (2004).

2. Sih, A., Bell, A. M. & Johnson, J. C. Behavioral syndromes: an integrative overview. The quarterly review of … (2004) doi:10.3233/jpd-120165.

3. Bergmüller, R. & Taborsky, M. Animal personality due to social niche specialisation. Trends in ecology & evolution 25, 504–511 (2010).

4. Kempermann, G. Environmental enrichment, new neurons and the neurobiology of individuality. Nature Reviews Neuroscience 78, 1–11 (2019).

5. Pennisi, E. The power of personality. 644–647 (2016).

6. Freund, J. et al. Emergence of Individuality in Genetically Identical Mice. Science (New York, N.Y.) 340, 756–759 (2013).

7. Torquet, N. et al. Social interactions impact on the dopaminergic system and drive individuality. Nature Communications 9, 3081 (2018).

8. Hayden, B. Y. Economic choice: the foraging perspective. Curr Opin Behav Sci 24, 1–6 (2018).

9. Hayden, B. Y. & Walton, M. E. Neuroscience of foraging. Frontiers in neuroscience 8, 81 (2014).

10. Mobbs, D., Trimmer, P. C., Blumstein, D. T. & Dayan, P. Foraging for foundations in decision neuroscience: insights from ethology. Nat Rev Neurosci 19, 419–427 (2018).

11. Timberlake, W. & Peden, B. F. On the distinction between open and closed economies. Journal of the experimental analysis of behavior 48, 35–60 (1987).

12. Rowland, N. E., Vaughan, C. H., Mathes, C. M. & Mitra, A. Feeding behavior, obesity, and neuroeconomics. Physiology & Behavior 93, 97–109 (2008).

13. Beeler, J. A., Daw, N., Frazier, C. R. M. & Zhuang, X. Tonic dopamine modulates exploitation of reward learning. Frontiers in Behavioral Neuroscience 4, 170 (2010).

14. Gartland, L. A., Firth, J. A., Laskowski, K. L., Jeanson, R. & Ioannou, C. C. Sociability as a personality trait in animals: methods, causes and consequences. Biol Rev (2021) doi:10.1111/brv.12823.

15. Skóra, M. N. et al. Personality driven alcohol and drug abuse: New mechanisms revealed. Neurosci. Biobehav. Rev. 116, 64–73 (2020).

16. Kreek, M. J., Nielsen, D. A., Butelman, E. R. & LaForge, K. S. Genetic influences on impulsivity, risk taking, stress responsivity and vulnerability to drug abuse and addiction. Nature Neuroscience 8, 1450– 1457 (2005).

17. Diergaarde, L. et al. Impulsive choice and impulsive action predict vulnerability to distinct stages of nicotine seeking in rats. Biological psychiatry 63, 301–308 (2008).

18. Perkins, K. A. et al. Initial nicotine sensitivity in humans as a function of impulsivity. Psychopharmacology 200, 529–544 (2008).

19. Perkins, K. A., Gerlach, D., Broge, M., Grobe, J. E. & Wilson, A. Greater sensitivity to subjective effects of nicotine in nonsmokers high in sensation seeking. Exp. Clin. Psychopharmacol. 8, 462–471 (2000).

20. Faure, P., Fayad, S. L., Solié, C. & Reynolds, L. M. Social Determinants of Inter-Individual Variability and Vulnerability: The Role of Dopamine. Front Behav Neurosci 16, 836343 (2022).

21. Addicott, M. A., Pearson, J. M., Sweitzer, M. M., Barack, D. L. & Platt, M. L. A Primer on Foraging and the Explore/Exploit Trade-Off for Psychiatry Research. Neuropsychopharmacology : official publication of the American College of Neuropsychopharmacology 42, 1931–1939 (2017).

22. Berke, J. D. What does dopamine mean? Nat Neurosci 21, 787–793 (2018).

23. Schultz, W. Multiple dopamine functions at different time courses. Annual review of neuroscience 30, 259–288 (2007).

24. Solié, C., Girard, B., Righetti, B., Tapparel, M. & Bellone, C. VTA dopamine neuron activity encodes social interaction and promotes reinforcement learning through social prediction error. Nat Neurosci 1– 12 (2021) doi:10.1038/s41593-021-00972-9.

25. Exley, R. et al. Distinct contributions of nicotinic acetylcholine receptor subunit alpha4 and subunit alpha6 to the reinforcing effects of nicotine. Proceedings of the National Academy of Sciences of the United States of America 108, 7577–7582 (2011).

26. Maskos, U. et al. Nicotine reinforcement and cognition restored by targeted expression of nicotinic receptors. Nature 436, 103–107 (2005).

27. Mameli-Engvall, M. et al. Hierarchical control of dopamine neuron-firing patterns by nicotinic receptors. Neuron 50, 911–921 (2006).

28. Morel, C. et al. Nicotine consumption is regulated by a human polymorphism in dopamine neurons. Molecular Psychiatry 19, 930–936 (2014).

29. Faure, P., Tolu, S., Valverde, S. & Naudé, J. Role of nicotinic acetylcholine receptors in regulating dopamine neuron activity. Neuroscience 282C, 86–100 (2014).

30. Naudé, J., Dongelmans, M. & Faure, P. Nicotinic alteration of decision-making. Neuropharmacology 96, 244–254 (2015).

31. Maubourguet, N., Lesne, A., Changeux, J.-P., Maskos, U. & Faure, P. Behavioral sequence analysis reveals a novel role for beta2* nicotinic receptors in exploration. PLoS Computational Biology 4, e1000229 (2008).

32. Addicott, M. A., Pearson, J. M., Wilson, J., Platt, M. L. & McClernon, F. J. Smoking and the bandit: A preliminary study of smoker and nonsmoker differences in exploratory behavior measured with a multiarmed bandit task. Experimental and Clinical Psychopharmacology 21, 66–73 (2013).

33. Sidorenko, N., et al. Acetylcholine and noradrenaline enhance foraging optimality in humans. Proc. Natl. Acad. Sci. 120, e2305596120 (2023).

34. Dongelmans, M. et al. Chronic nicotine increases midbrain dopamine neuron activity and biases individual strategies towards reduced exploration in mice. Nat Commun 12, 6945 (2021).

35. Barik, J. et al. Chronic stress triggers social aversion via glucocorticoid receptor in dopaminoceptive neurons. Science (New York, N.Y.) 339, 332–335 (2013).

36. Chaudhury, D. et al. Rapid regulation of depression-related behaviours by control of midbrain dopamine neurons. Nature 493, 532–536 (2013).

37. Morel, C. et al. Nicotinic receptors mediate stress-nicotine detrimental interplay via dopamine cells’ activity. Molecular Psychiatry 23, 1597–1605 (2017).

38. Pomerleau, O. F. Individual differences in sensitivity to nicotine: Implications for genetic research on nicotine dependence. Behav Genet 25, 161–177 (1995).

39. Mondoloni, S., et al. Prolonged nicotine exposure reduces aversion to the drug in mice by altering nicotinic transmission in the interpeduncular nucleus. Biorxiv 2021.12.16.472949 (2022) doi:10.1101/2021.12.16.472949.

40. Faure, P., Neumeister, H., Faber, D. S. & Korn, H. Symbolic analysis of swimming trajectories reveals scale invariance and provides a model for fish locomotion. World Scientific 11, 233–243 (2003).

41. Sutton, R. S. & Barto, A. G. Reinforcement Learning. (MIT Press, 1998).

42. Rutledge, R. B. et al. Dopaminergic drugs modulate learning rates and perseveration in Parkinson’s patients in a dynamic foraging task. The Journal of neuroscience : the official journal of the Society for Neuroscience 29, 15104–15114 (2009).

43. Renier, N. et al. Mapping of Brain Activity by Automated Volume Analysis of Immediate Early Genes. Cell 165, 1789–1802 (2016).

44. Pich, E. M., Chiamulera, C. & Tessari, M. Neural substrate of nicotine addiction as defined by functional brain maps of gene expression. J. Physiol.-Paris 92, 225–228 (1998).

45. Pagliusi, S. R., Tessari, M., DeVevey, S., Chiamulera, C. & Pich, E. M. The Reinforcing Properties of Nicotine are Associated with a Specific Patterning of c-fos Expression in the Rat Brain. Eur. J. Neurosci. 8, 2247–2256 (1996).

46. Eddine, R. et al. A concurrent excitation and inhibition of dopaminergic subpopulations in response to nicotine. Scientific Reports 5, 8184 (2015).

47. Nguyen, C. et al. Nicotine inhibits the VTA-to-amygdala dopamine pathway to promote anxiety. Neuron (2021) doi:10.1016/j.neuron.2021.06.013.

48. Gomez-Marin, A. & Ghazanfar, A. A. The Life of Behavior. Neuron 104, 25–36 (2019).

49. Lopes, J. B. et al. Individual behavioral trajectories shape whole-brain connectivity in mice. Biorxiv 2022.03.25.485806 (2022) doi:10.1101/2022.03.25.485806.

50. Markowitz, J. E. et al. Spontaneous behaviour is structured by reinforcement without explicit reward. Nature 1–10 (2023) doi:10.1038/s41586-022-05611-2.

51. Coddington, L. T., Lindo, S. E. & Dudman, J. T. Mesolimbic dopamine adapts the rate of learning from action. Nature 1–9 (2023) doi:10.1038/s41586-022-05614-z.

52. Dember, W. N. & Fowler, H. Spontaneous alternation behavior. Psychological Bulletin 55, 412–428 (1958).

53. Sandi, C. Understanding the neurobiological basis of behavior: a good way to go. Frontiers in neuroscience 2, 129–130 (2008).

54. Schaefer, A. T. & Claridge-Chang, A. The surveillance state of behavioral automation. Current opinion in neurobiology 22, 170–176 (2012).

55. Spruijt, B. M. & DeVisser, L. Advanced behavioural screening: automated home cage ethology. Drug Discovery Today: Technologies 3, 231–237 (2006).

56. Castelhano-Carlos, M., Costa, P. S., Russig, H. & Sousa, N. PhenoWorld: a new paradigm to screen rodent behavior. Translational Psychiatry 4, e399–11 (2014).

57. Ungless, M. A. & Grace, A. A. Are you or aren’t you? Challenges associated with physiologically identifying dopamine neurons. Trends in Neurosciences 35, 422–430 (2012).

58. Renier, N., et al. iDISCO: A Simple, Rapid Method to Immunolabel Large Tissue Samples for Volume Imaging. Cell 159, 896–910 (2014).

59. Klein, S., Staring, M., Murphy, K., Viergever, M. A. & Pluim, J. P. W. elastix: a toolbox for intensity-based medical image registration. IEEE Trans. Méd. imaging 29, 196–205 (2009).

60. Wang, Q. et al. The Allen Mouse Brain Common Coordinate Framework: A 3D Reference Atlas. Cell 181, 936–953.e20 (2020).

61. Daw, N. D. Trial-by-trial data analysis using computational models. in Decision Making, Affect, and Learning 3–38 (Oxford University Press, 2011). doi:10.1093/acprof:oso/9780199600434.003.0001.

62. Eugster, M. J. A. & Leisch, F. From Spider-Man to Hero - Archetypal Analysis in R. Journal of Statistical Software 30, 1–23 (2009).

